# Two distinct binding modes provide the RNA binding protein RbFox with extraordinary sequence specificity

**DOI:** 10.1101/2021.12.23.474020

**Authors:** Xuan Ye, Wen Yang, Soon Yi, Yanan Zhao, Fan Yang, Gabriele Varani, Eckhard Jankowsky

## Abstract

The specificity of RNA-binding proteins for their target sequences varies considerably. Yet, it is not understood how certain proteins achieve markedly higher sequence specificity than most others. Here we show that the RNA Recognition Motif of RbFox accomplishes extraordinary sequence specificity by employing functionally and structurally distinct binding modes. Affinity measurements of RbFox for all binding site variants reveal the existence of two different binding modes. The first exclusively binds the cognate and a closely related RNA variant with high affinity. The second mode accommodates all other RNAs with greatly reduced affinity, thereby imposing large thermodynamic penalties on even near-cognate sequences. NMR studies indicate marked structural differences between the two binding modes, including large conformational rearrangements distant from the RNA binding site. Distinct binding modes by a single RNA binding module explain extraordinary sequence selectivity and reveal an unknown layer of functional diversity, cross talk and regulation for RNA-protein interactions.

## INTRODUCTION

In eukaryotic cells, the expression of tens of thousands of RNAs is regulated by thousands of diverse RNA binding proteins (RBPs)^1,2^. A pivotal aspect of this regulation is the specificity of RBPs for their target sites^1–5^; that is, the degree by which an RBP discriminates between cognate and non-cognate RNAs^1,2,4^. Specificity of RBPs varies considerably; some RBPs are highly selective for certain RNA sites, others bind degenerate regions, and yet other RBPs interact with RNAs broadly, with little sequence discrimination^6–12^. Defining the molecular mechanisms through which RBPs acquire their specificity is critical for understanding the rules of RNA biology and for devising therapeutic strategies against diseases associated with RBP- and RNA-related processes^1,2,13^.

Among RBPs with the greatest specificity are the three closely related human RbFox proteins (RbFox 1-3), which function in pre-mRNA splicing, miRNA processing and other RNA metabolic steps^14^. The three RbFox proteins contain divergent N- and C-terminal regions that are likely unstructured, but share a highly conserved RNA binding domain that belongs to the <u>R</u>NA <u>R</u>ecognition <u>M</u>otif (RRM) superfamily^14,15^. The RbFox RRM binds RNAs with a consensus 5’-GCAUG motif with low nanomolar affinity *in vitro*^16–19^. Even minor changes in the consensus motif decrease the RNA affinity by orders of magnitude^17,19^, indicating an inherent selectivity of RbFox proteins for their consensus sequence that is markedly higher than for most other RBPs^6,8,9^. In the cell, however, biological effects by RbFox proteins are also exerted by binding to non-consensus sites, provided expression levels of RbFox increase^20^.

How RbFox proteins accomplish higher specificity than most other RBPs is not understood. Even the existing NMR structures, which provide detailed views of the RNA binding interface, fail to provide a cogent reason for this extraordinary specificity^17,18^. We therefore set out to systematically examine the basis for the high specificity of the RbFox proteins through a combination of high throughput biochemical techniques followed by NMR structure determination. Simultaneous affinity measurements of the RRM of RbFox for all possible 16,348 7-mer RNA sequence variants revealed two distinct binding modes, one associated with binding to the consensus 5’-GCAUG and a second mode for binding to all other sequences. Mutations in RbFox only have a small effect on the binding of the consensus sequence, but markedly increase affinity for all other variants, thereby diminishing specificity. The results indicate that the binding mode for the non-consensus sequences enables the imposition of large thermodynamic penalties even on near-cognate variants, thereby accomplishing exquisite discrimination between consensus and non-consensus RNAs. Comparison of the NMR structures with a non-consensus and the consensus sequence reveals that the two binding modes are associated with substantial structural differences within the RRM. Most remarkably, the structural rearrangement extends to protein regions distant from the RNA binding interface, suggesting that the distinct binding modes can transmit RNA sequence information to potential protein binding partners of RbFox.

Thus, our data show that RbFox employs a previously unknown mechanism to accomplish its extraordinary specificity - structurally distinct binding modes that enable the imposition of large thermodynamic penalties on non-consensus sequences and transmit RNA sequence information to distant regions of the RRM. The results reveal a novel layer of biological complexity in RNA-protein interactions and post-transcriptional regulation of gene expression.

## RESULTS

### The affinity distribution of RbFox is bimodal

To examine how RbFox achieves high specificity, we first measured the affinity of the RNA binding domain (RRM) of RbFox for two RNAs that differ by a single nucleotide (**Fig.1a**). The apparent equilibrium binding constant (*K*_1/2_) for the RbFox cognate sequence (5’-GCAUG) was reduced by orders of magnitude by this single nucleotide substitution (**Fig.1a,b**), consistent with previous data^17–19^. This result highlights the remarkable ability of RbFox to discriminate effectively between RNA substrates that differ by only small sequence variations.

**Figure 1.**
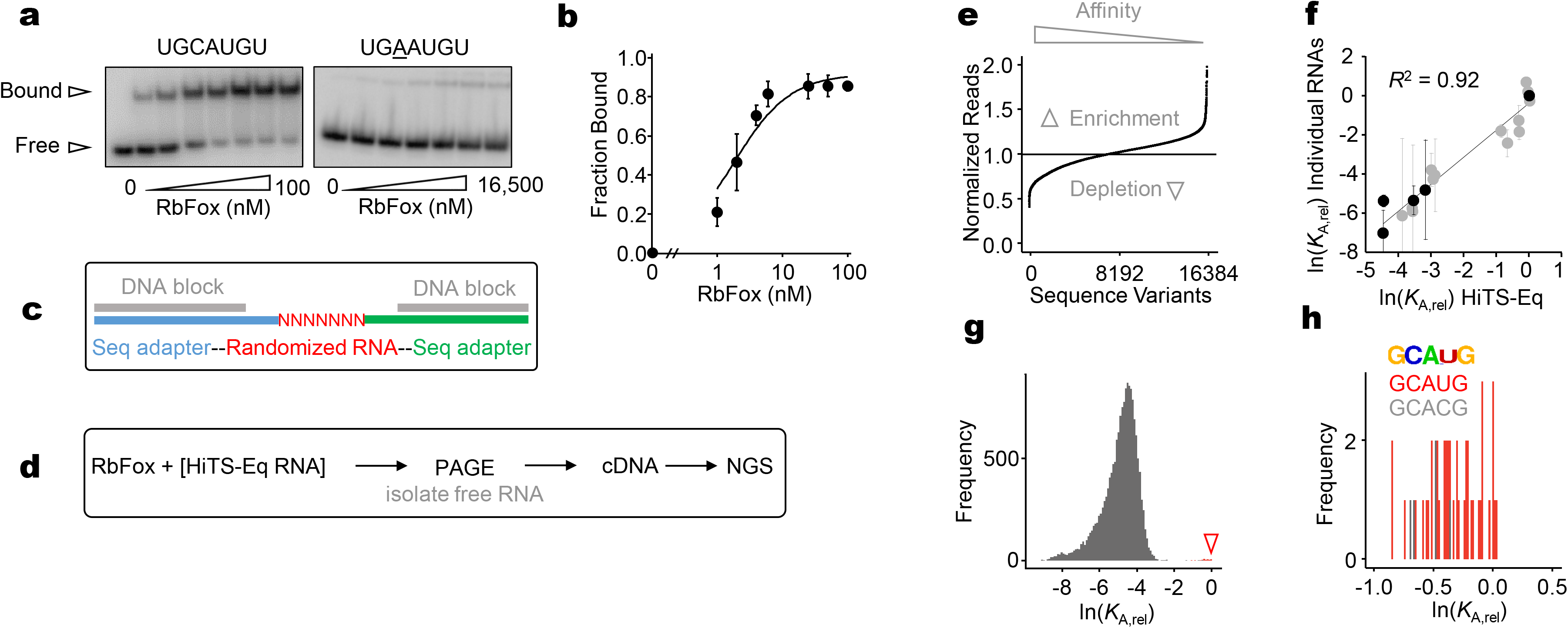
RbFox binding to all 7-mer RNA sequence variants. **a**, Representative PAGE images for RbFox binding to individual RNA substrates under equilibrium conditions.(RbFox: 0, 1, 2, 4, 6, 25, 50 and 100 nM for 5’-U<u>GCAUG</u>U, underline marks the consensus 5-mer; 258, 516, 1,031, 2,063, 4,125, 8,250 and 16,500 nM for 5’-U<u>GAAUG</u>U, RNAs: 1 nM). **b**, Binding isotherm for 5’-U<u>GCAUG</u>U (data points: average of three independent experiments; error bars: one standard deviation; line: best fit to binding isotherm with *K*_1/2_^(U<u>GCAUG</u>U)^ = 1.6 ± 0.3 nM). The low level of RbFox binding to 5’-U<u>GAAUG</u>U (panel **a**) precludes reliable affinity determination (estimated lower limit for *K’*_1/2_(U<u>GAAUG</u>U) > 17 μM). **c**, Design of RNA substrate pool for the HiTS-Eq measurements (detailed information: **Suppl. Fig.S1**). **d**, Basic Scheme for the HiTS-Eq approach. **e**, Depletion (normalized reads < 1) and enrichment (normalized reads > 1) of RNA sequence variants at [RbFox] = 19.74 μM. Reads are normalized to read numbers in the library without protein. **f**, Relative apparent association constants (*K*_A,rel_) for corresponding RNA variants (for variants, see Materials and Methods), measured for individual RNA and by HiTS-Eq (data points: average of three independent experiments; error bars: one standard deviation; *R^2^*: correlation coefficient; black points: our measurements; grey points: values reported by Stoltz *et al*^19^). **g**, Affinity distribution (*K*_A,rel_) of RbFox for all 16,384 RNA sequence variants (bin size: 100). The triangle indicates the population of high affinity variants. **h**, Distribution of high affinity variants (bin size: 100). (Sequence motif logo: determined for 40 variants with the highest affinity^48,49^; E = 2.7e^−77^; red bins: variants containing 5’-GCAUG; grey bins: variants containing 5’-GCACG).

Next, we comprehensively characterized the specificity landscape of the RbFox RRM by employing <u>Hi</u>gh <u>T</u>hroughput <u>S</u>equencing <u>Eq</u>uilibrium Binding (HiTS-Eq)^10,11,21,22^. This approach allows the simultaneous determination of apparent affinities for large numbers of sequence variants within an RNA pool^10,11,21^. We constructed a pool of 16,384 unique RNAs encoding all possible 7-mer sequences in a segment with 7 randomized nucleotides (**Fig.1c**). This segment was flanked on each side by a 3 nt linker with fixed sequence and an adaptor to facilitate conversion of the RNA into cDNA for subsequent Illumina sequencing (**Fig.1c, Extended Data Fig.1**). To prevent base pairing between adapter regions and complementary sequences in the randomized RNA regions, which would bias the available substrate pool^21,22^, we annealed DNAs complementary to the adapter regions to mask the corresponding sequences (**Fig.1c**).

The RNA pool was incubated in separate reactions with increasing concentrations of RbFox. Upon reaching equilibrium, samples were transferred to non-denaturing PAGE to separate RbFox-bound and -unbound substrates (**Extended Data Fig.1**). Unbound substrates were isolated, converted to cDNA, and sequenced on the Illumina platform (**Fig.1d**). The preparation of the cDNA libraries included unique molecular identifiers (UMIs), to identify and correct for PCR overamplification artifacts^7,21,22^ (**Extended Data Fig.1**). As expected, the addition of RbFox caused depletion of variants in the unbound RNA pool, compared to the control pool without RbFox (**Fig.1e**).

We calculated apparent relative association constants (*K*_A(rel)_, affinities normalized to the affinity of the sequence 5’-UGCAUGU) for all substrate variants from binding reactions with increasing RbFox concentrations, as previously described^17–19^. Experiments were performed in independent duplicates, which provided highly correlated sequencing read values (*R*^2^ > 0.85; **Extended Data Fig.2**). Apparent affinities obtained with HiTS-Eq also correlate well with affinities measured for individual substrates by us and others^19^ (*R*^2^ = 0.92, **Fig.1f**), indicating that the HiTS-Eq approach faithfully reflects results of conventional biochemical assays.

The histogram of *K*_A(rel)_ values for all sequence variants represents the global affinity distribution for RbFox (**Fig.1g**). Notably, the distribution is bimodal, with a large, broad peak encompassing the vast majority of the 16,384 sequence variants, and a much smaller peak with sequence variants that contain the cognate 5’-GCAUG and the closely related 5’-GCACG (**Fig.1g**, small peak marked by triangle). High affinities of RbFox for 5’-GCACG containing variants are consistent with previous observations^6,9,16^. Accordingly, the sequence logo for sequence variants with the highest affinities is identical to that obtained with the RNA Bind-n-Seq (RBNS), RNAcompete and HTR-SELEX approaches^6,8,9,16^ (**Fig.1h**).

Our data provide two new insights. First, the affinity of RbFox for all sequence variants without 5’-GCA(U/C)G is lower by orders of magnitude, compared to its cognate 5-mer. To our knowledge, this extraordinary ability to discriminate against all but two closely related sequence variants is unparalleled by other RBPs. Second, the bimodal affinity distribution of RbFox differs from the unimodal distributions seen for other RBPs^6,7,10,11^.

### Quantitative binding models indicate two distinct binding modes

To understand the determinants of these unique features of RbFox specificity, we applied quantitative binding models, focusing on the minimal 5 nt RbFox consensus^7,10,22^. To account for variations in the register of each possible 5-mers in our RNA pool with 7 randomized nucleotides, we plotted affinity values for all 48 5-mers for each of the 1,024 5-mer variants and determined the median relative affinity for each 5-mer (**Fig.2a**, **Extended Data Fig.3a**). The obtained values correlate well with corresponding 5-mer affinity scores calculated from the RBNS approach (**Extended Data Fig.3b**), a high throughput approach designed to delineate preferred binding motifs for RBPs^6,16^.

**Figure 2.**
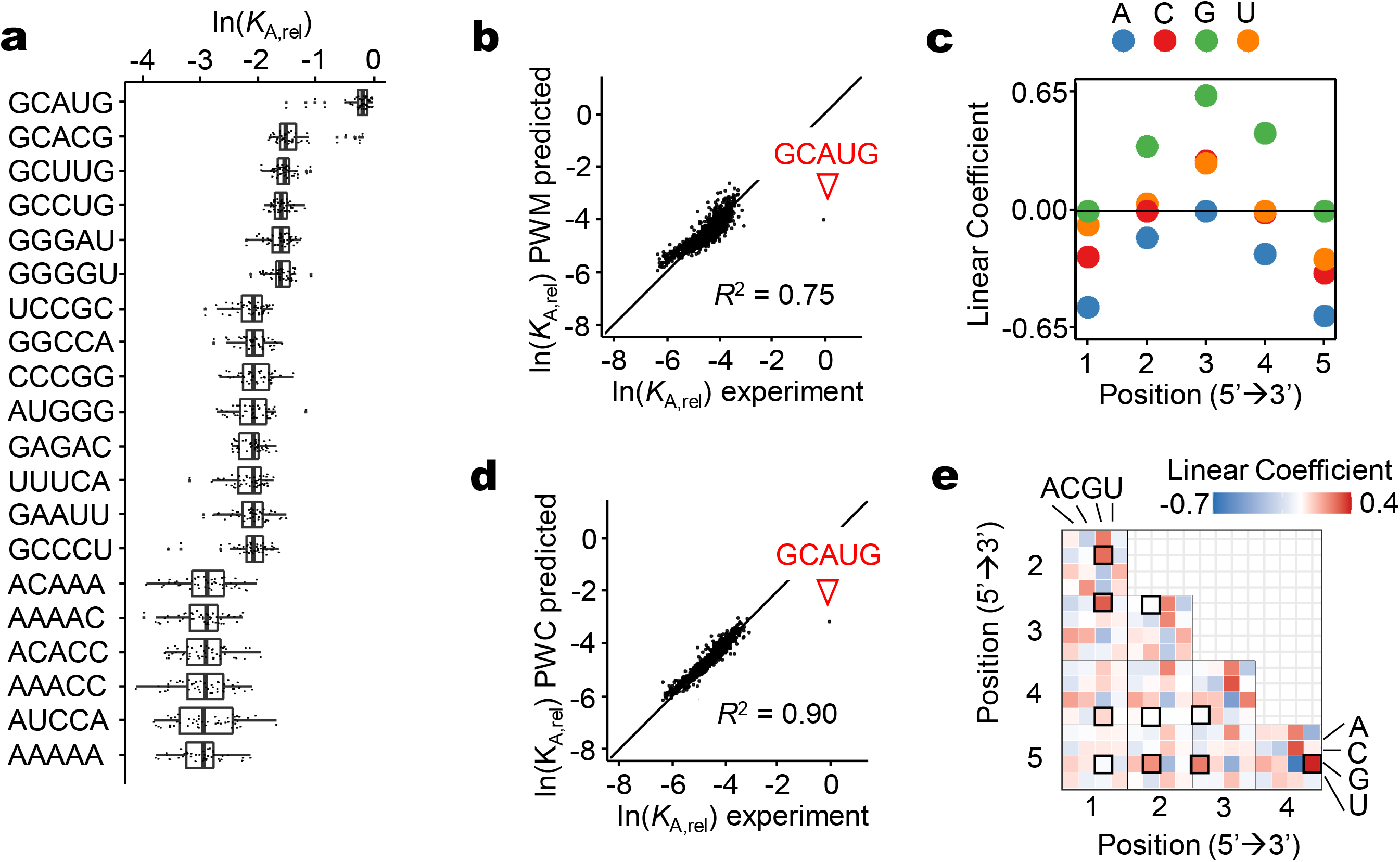
Analysis of RbFox affinity distribution with quantitative binding models. **a**, Relative affinities (*K*_A,rel_) for selected 5-mer RNA variants, indicated on the right. 48 *K*_A,rel_ values correspond to each 5-mer (vertical line: median; box: variability through lower quartile and upper quartile; whiskers: variability outside the lower and upper quartiles). **b**, Correlation between experimental *K*_A,rel_ values for each 5-mer (median value, panel **a**) and values calculated with the Position Weight Matrix (PWM) binding model (triangle: consensus 5-mer; line: diagonal, y = x; *R^2^*: correlation coefficient). **c**, Linear coefficients for each nucleotide position calculated with the PWM binding model (negative values: destabilization). **d**, Correlation between experimental *K*_A,rel_ values for each 5-mer (median value, panel **a**) with values calculated with the Pairwise Coupling (PWC) binding model (triangle: consensus 5-mer; line: diagonal, y = x; *R^2^*: correlation coefficient). **e**, Linear coefficients for each pairwise coupling between all nucleotides calculated with the PWC binding model (black frames: couplings in consensus 5-mer).

Comparison of the HiTS-Eq 5-mer and 7-mer affinities show how nucleotides flanking the core 5-mer affect RbFox binding (**Extended Data Fig.3c**). Comparable affinities of 5’-GCAUG and 5’-GCACG variants depend on nucleotides flanking the core 5-mer, as previously noted^6,9,16^. Our data reveal that the impact of flanking nucleotides depends also on the 5-mer sequence. For example, a 5’-U flanking 5’-GCACG enhances the affinity, compared to a 5’-G, but for most other 5-mers, a 5’-U decreases the affinity, compared to 5’-G (**Extended Data Fig.3c**).

To better understand the rules governing binding of RbFox to the core 5-mer, we analyzed the median affinity values with a Position Weight Matrix (PWM) model (**Fig.2b**). Although a PWM considers only the position of each nucleotide in isolation^7,10,22^, the model accounts for 75% of the data variance among the 5-mers (**Fig.2b**). However, the PWM bears no resemblance to the cognate 5-mer, indicating instead a preference for G at all positions (**Fig.2c**). Most interestingly, the model reveals a single outlier – the cognate 5’-GCAUG sequence (**Fig.2b**). These observations indicate that a PWM can describe affinities for almost all sequence variants to a considerable degree, except for the cognate variant.

To examine whether the inherent limitations of the PWM cause the striking divergence between cognate and non-cognate RNAs, we applied a binding model that considers pairwise coupling (PWC) between all nucleotides^7,10,22,23^. The PWC model accounts for 90% of the data variance (**Fig.2d**), an outstanding result for experimental high-throughput data^7,10,11^. Yet, the cognate 5-mer remains a clear outlier, even though the PWC model highlights favorable base coupling contributions resembling the cognate variant (**Fig.2e**, highlighted fields). PWM and PWC models for all 7-mer RNA variants, which do not consider the binding register of the minimal 5-mer, also identify variants with the cognate 5’-GCAUG as outliers (**Extended Data Fig.4**). Collectively, our analyses of RbFox binding with quantitative binding models suggests that RbFox employs distinct binding modes – one for its cognate 5-mer and one and a second mode for non-cognate sequence variants. To our knowledge, multiple binding modes have never been reported for a single RRM.

### Protein mutations affect the two binding modes differently

To further probe the notion of two distinct binding modes, we measured the affinity distribution of an RbFox variant containing four amino acid changes (RbFox^mut^, **Fig.3a**). These mutations were introduced to improve binding to a pri-miRNA comprising a near-consensus sequence^18^. We hypothesized that mutations in the RRM should affect the two distinct binding modes differently. We first measured binding of RbFox^mut^ to the individual RNAs examined above with wild type (wt) RbFox (**Fig.3b**). RbFox^mut^ bound an RNA with the cognate 5-mer with slightly reduced affinity, compared to wt RbFox (**Fig.3b,c**). However, RbFox^mut^ bound to an RNA with a single base change only slightly less well than to the cognate sequence. In contrast, binding of wt RbFox to this RNA was barely detectable (**Fig.1a,b**). This observation indicates that the mutations in RbFox^mut^ markedly increase the affinity for non-cognate RNAs, compared to the wt protein.

**Figure 3.**
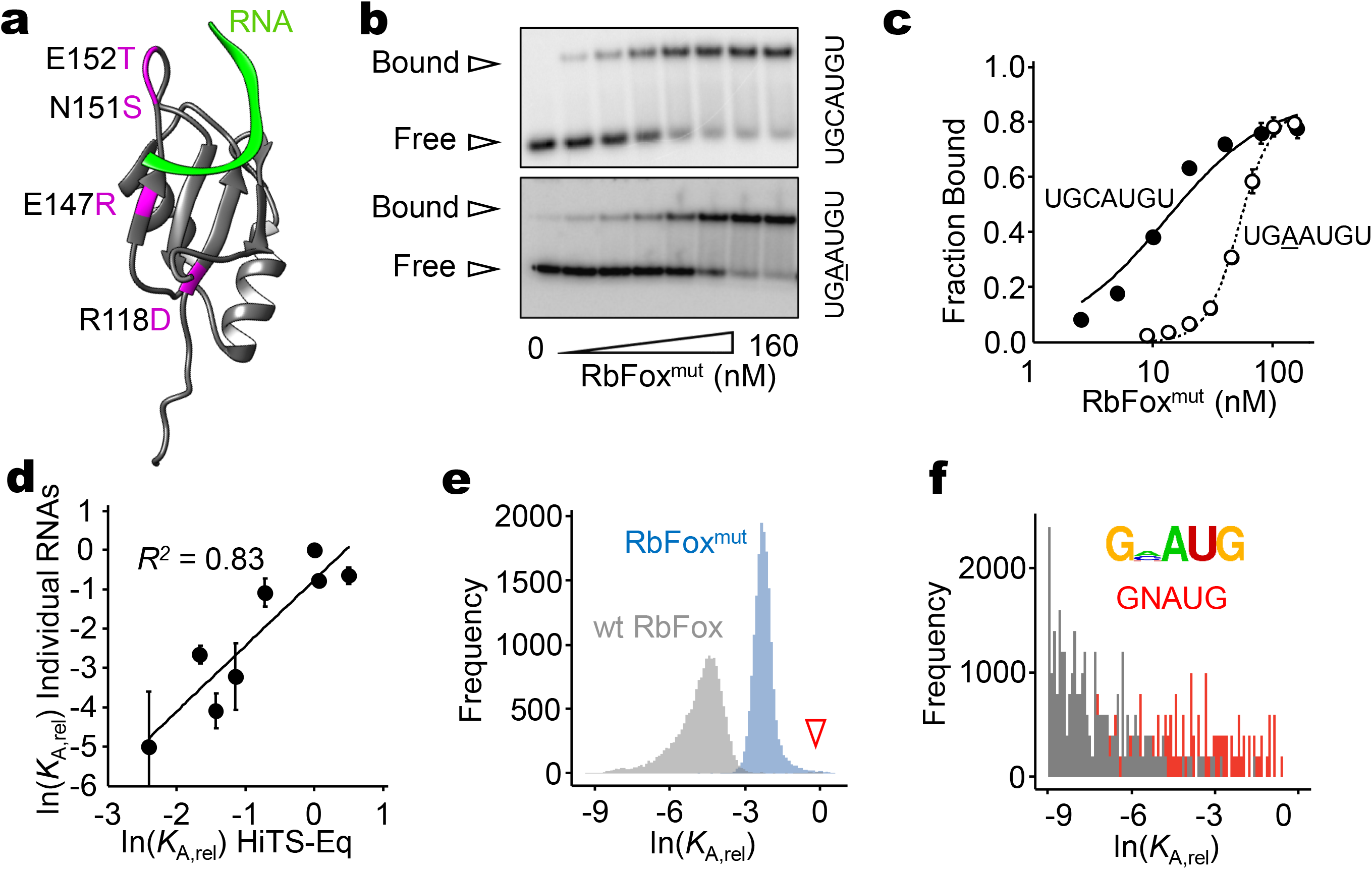
Impact of RbFox mutations on the affinity distribution. **a**, Locations of the mutations in RbFox^mut^ in the RRM^18^, highlighted in purple (green band: RNA: 5’-U<u>GCAUG</u>U^17^). **b**, Representative PAGE images for RbFox^mut^ equilibrium binding to individual RNA substrates (sequences on the right; RbFox^mut^: 0, 2.5, 5, 10, 20, 40, 80 and 160 nM for both substrates: RNAs: 1 nM). **c**, Equilibrium binding isotherm for RbFox^mut^ with the substrates shown in panel **a**(data points: average of three independent experiments; error bars: one standard deviation; lines: best fit to binding isotherm; *K*_1/2_(U<u>GCAUG</u>U) = 14.3 ± 2.5 nM, *K*_1/2_(U<u>GAAUG</u>U) = 35.3 ± 11.8 nM). **d**, Relative apparent association constants (*K*_A,rel_) for corresponding RNA variants (for sequences, see Materials and Methods), measured for individual RNAs and by HiTS-Eq (data points: average of three independent experiments; error bars: one standard deviation; *R^2^*: correlation coefficient). **e**, Affinity distribution (*K*_A,rel_) of RbFox^mut^ for all 7-mer RNA sequence variants (blue). (bin size: 100). For reference, the affinity distribution of wild type RbFox (grey) is plotted as well (triangle: population of high affinity variants for wt RbFox). **f**, Distribution of high affinity variants for RbFox^mut^ (bin size: 100). (Sequence motif logo was determined for 40 variants with the highest affinity^48,49^; E = 5.6e^−57^; red bins: variants containing 5’-GNAUG; grey bins: variants that differ from 5’-GNAUG).

We next examined RbFox^mut^ with HiTS-Eq using the RNA pool employed above for wt RbFox (**Extended Data Fig.5**). Apparent affinities obtained with HiTS-Eq correlate well with corresponding data measured for individual RNAs (*R*^2^ = 0.83, **Fig.3d**). However, the affinity distribution for RbFox^mut^ does not display the pronounced bimodal shape observed with wt RbFox. Rather, the RbFox^mut^ distribution shows a tail corresponding to high affinity variants (**Fig.3e, Extended Data Fig.6a**). In addition, the RbFox^mut^ distribution narrows and is shifted towards higher relative affinities compared to the wt RbFox distribution (**Fig.3e**). The global increase in relative affinities for non-cognate sequences, compared to wt RbFox, coincides with a pronounced decrease in specificity for RbFox^mut^ (**Fig.3e, Extended Data Fig.6a**); that is, the mutant protein does not discriminate against non-cognate sequences as strongly as the wt protein. In addition, the base preference of wt RbFox at position 2 of the cognate 5-mer is essentially lost in RbFox^mut^ (**Fig.3f, Fig1h**); a possible reason for the slightly lower affinity of RbFox^mut^ for the cognate sequence, compared to wt RbFox (**Fig.3b**). The loss of stringency at position 2 also allows RbFox^mut^ to accommodate more diverse 5-mer variants in the binding mode reserved for the cognate sequence in wt RbFox.

The opposite effects of RbFox mutations on the affinities for cognate and non-cognate RNAs, strongly supports the notion of distinct binding modes of RbFox. In addition, our data reveal that wt RbFox accomplishes high specificity by thermodynamically penalizing all sequences, except the cognate variant and the near cognate variant with a specific flanking nucleotide. RbFox^mut^ is unable to impose a similar thermodynamic penalty. The affinities for non-cognate sequence variants increase relative to the cognate sequence and specificity is reduced, compared to wt RbFox. The apparent need to impose a thermodynamic penalty on all but the cognate sequence provides an additional rationale for the two distinct binding modes observed for wt RbFox (**Fig.2**).

We next probed how the binding mode for the non-cognate RNA variants differed between wt and mutant RbFox. To this end we analyzed the affinity distribution of RbFox^mut^ with the PWM and the PWC models, as described for wt RbFox (**Extended Data Fig.6b**). The PWM model for RbFox^mut^ accounted for 35% of the data variance and differed from the PWM for wt RbFox, most notably by a preference for A, instead of G (**Extended Data Fig.7**), a feature that was successfully designed for in the mutant protein^18^. The PWC model accounted for 66% of the data variance for RbFox^mut^ (**Fig.4a,b**) and revealed numerous differences with wt RbFox (**Fig.4c**). Collectively, these data indicate substantial differences in the binding mode for the non-cognate variants between RbFox^mut^ and wt protein, thereby supporting the notion of distinct binding modes.

**Figure 4.**
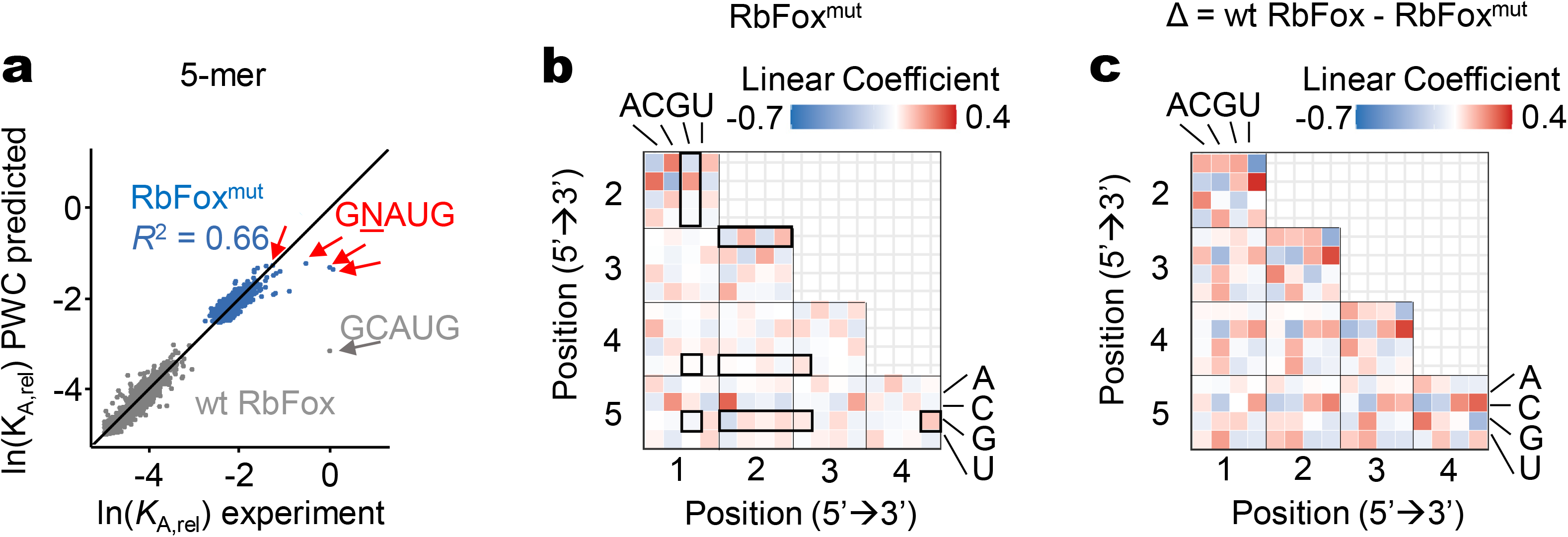
Analysis of RbFox^mut^ affinity distribution with quantitative binding models. **a**, Correlation between experimental *K*_A,rel_ values for RbFox^mut^ for each 5-mer (median values, **Suppl. Fig.S6**) with values calculated with the PWC binding model (blue dots: RbFox^mut^; red arrow: consensus 5-mer 5’-GNAUG; line: diagonal, y = x; *R^2^*: correlation coefficient). The plot for wt RbFox (grey dots) is plotted as reference. **b**, Linear coefficients for each pairwise coupling between all nucleotides calculated with the PWC binding model; (black frames: couplings in consensus 5-mer, negative values: destabilization). **c**, Differences between linear coefficients for PWC binding model for wt RbFox, compared to RbFox^mut^; (positive values: increase in RbFox^mut^, compared to wt RbFox, negative values: decrease in RbFox^mut^, compared to wt RbFox).

### Cognate and non-cognate RNAs induce distinct structures in the RbFox RRM

The existence of two distinct binding modes of RbFox raised the question whether these binding modes also differ structurally. To examine this possibility, we performed NMR studies with a 7 nts cognate (5’-UGCAUGU) and non-cognate RNAs, which differs from the cognate variant at a single nucleotide (5’-UGCAU**<u>A</u>**U). We titrated each RNA into the wt RbFox RRM until saturation, while monitoring ^1^H-^15^N HSQC (**Extended Data Fig.8**). We observed slow-exchange binding kinetics with the cognate RNA and intermediate-exchange binding kinetics with the non-cognate variant (**Extended Data Fig.8**), consistent with the nanomolar affinity of the cognate RNA and the markedly lower dissociation constants of the non-cognate variant observed in the HiTS-Eq data (**Fig.2**).

Next, we collected 3D NMR spectra at saturating RNA concentrations and calculated the chemical shift difference between the two RbFox-RNA complexes (**Fig.5a,b**). Mapping of these differences onto the structure of wt RbFox identified changes in four regions of the protein: β1(R118-V121), β2 and the following loop (I143-E152), β3 (G159-E164) and the C-terminal tail (A191-A193) (**Fig.5c**). In the structure of RbFox RRM complexed with its cognate RNA, the AUG (nt 3 to 5) element is bound in a canonical manner through *π*-*π* stacking with the β-sheet surface, while UGC (nt −1 to 2) is recognized by loop residues^17^. Accordingly, similar chemical shifts for the region of RbFox interacting with UGC were observed in two complexes, indicating that the UGC nucleotides in both structures maintain the same contacts within the RNA binding cleft of RbFox. In contrast, the G-A substitution at position 5 causes marked changes in the β-sheet region that binds to the AUG element (0.2 - 0.7ppm, **Fig.5a**), as expected. Most notably, we also observed significant chemical shift changes (> 0.1ppm) in the β3/α2 and α1/β2 loops, which are distal to the RNA binding surface (**Fig.5a-c**).

**Figure 5.**
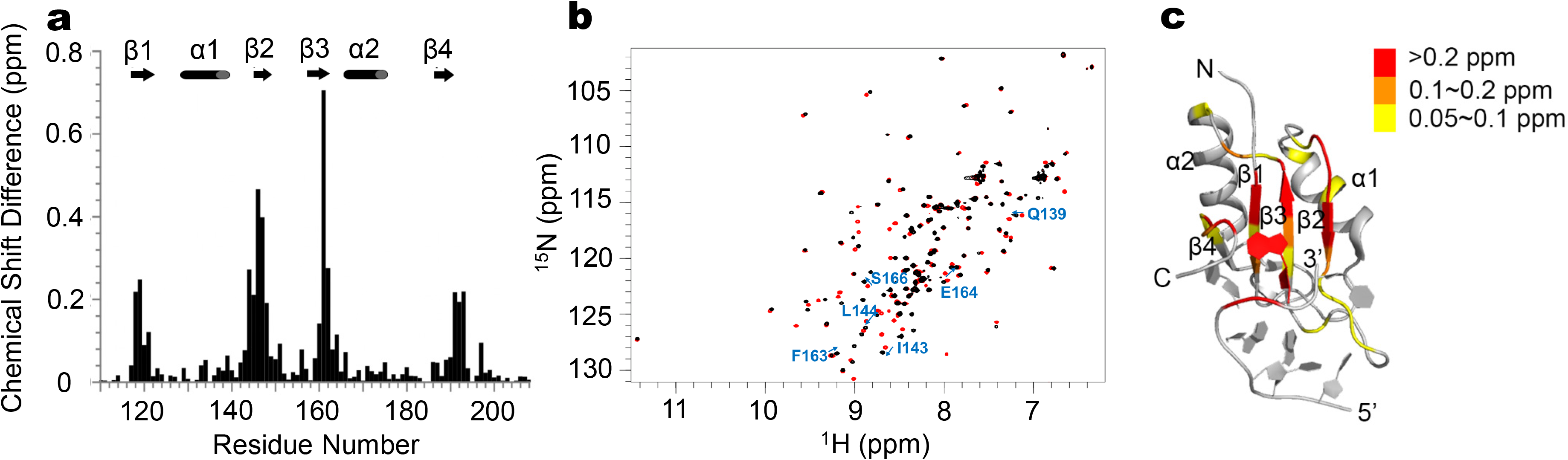
NMR analysis of the interaction of RbFox RRM with 5’-UGCAUGU and 5’-UGCAUAU. **a**, Chemical shift difference (CSD) of RbFox RRM bound to the two different RNA substrates. **b**, Superposition of ^1^H-^15^HSQC spectra of RbFox RRM complexed with the two RNAs (red: 5’-UGCAUGU; black: 5’-UGCAUAU). Residues with significant CSDs distant from the RNA binding site are labeled. **c**, Mapping of the CSD in panel **a** onto RbFox structure (red: CSD > 0.2 ppm; orange: 0.1 ppm < CSD < 0.2 ppm; yellow: 0.05 ppm < CSD < 0.1 ppm).

To gain more detailed insight into the structural differences between the RbFox RRM structures with cognate and non-cognate RNAs, we determined the structure of the RbFox RRM with the non-cognate RNA. The ^1^H-^15^N HSQC and NOESY spectra changed during data collection, suggesting multiple binding orientations that are most likely due to the weak binding of the non-cognate RNA. These characteristics made unambiguous NOE identification difficult. To overcome this problem, we borrowed NOEs from our previous NMR data for RbFox complexed to GCA within the miR-20b stem loop^24^. Since chemical shifts are very similar for both proteins and RNAs, we obtained a dataset that converged to a reliable structural ensemble (**Extended Data Table 1**).

A 2.0 Å rmsd difference was observed for RbFox in complex with the cognate and non-cognate RNAs, with large differences in protein regions distant to the RNA binding site as well (**Fig.6**). In the NMR structure with the cognate sequence, G6 forms the most extensive contacts with the protein (**Fig.6a**, left panel). In our structure with the non-cognate RNA, the G-A5 substitution abolishes the hydrogen-bonding interactions with R118 and T192. Our structure also reveals increased dynamics for A5 and the stacking interaction between the aromatic chain of F160 and the aliphatic chain of R194 (**Fig.6a**, right panel). The hydrogen bonding network of U4 is also rearranged due to loosened interactions between A5 and the protein. With the non-cognate RNA, the stacking between the imidazole ring of H120 and U4 is lost or weakened, because the side chain of H120 is flexible, as revealed by line broadening of the NMR spectra. The hydrogen-bonding interactions with N190 are completely lost, and those with T192 can only be observed for less than half of the models in the structural ensemble (**Fig.6a**). For a large majority of the converged structures, the aromatic ring of F160 tilts to stack with the base of A5. The RNA binding strands β1 and β3 float away from the RNA, due to the loss of close contacts with A5 and U4, and the β2 β3 loop reorients to better accommodate the non-cognate RNA (**Fig.6b**).

**Figure 6.**
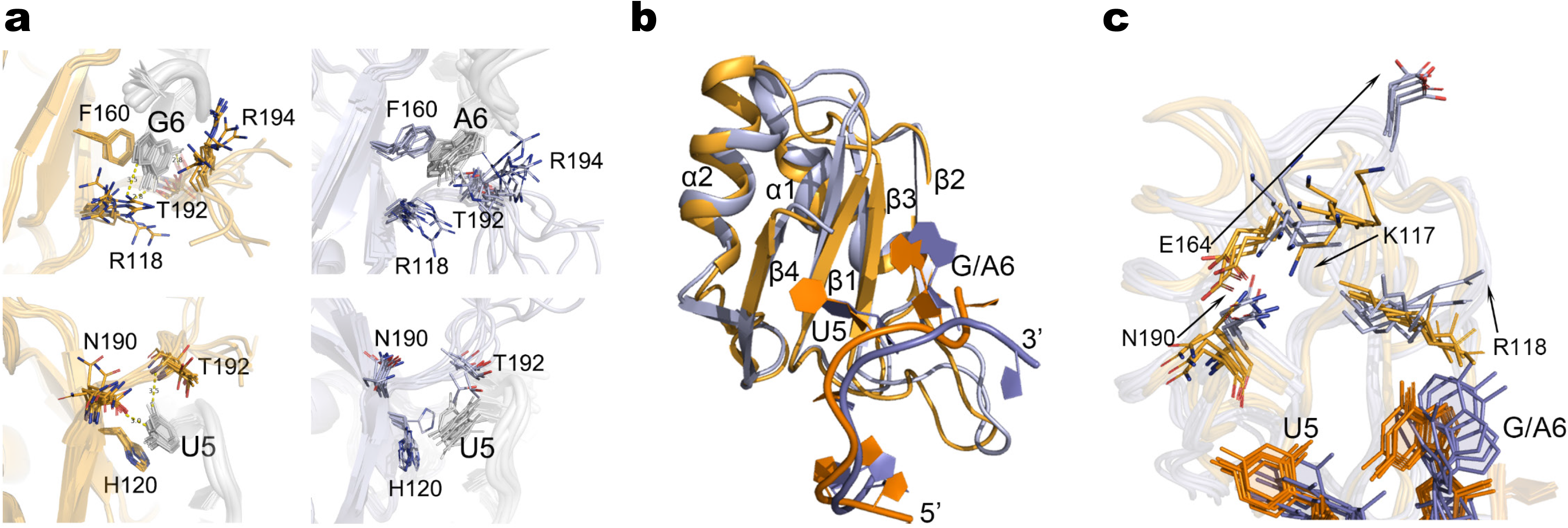
Comparison of RbFox RRM structures with 5’-UGCAUGU and 5’-UGCAUAU. **a**, Recognition of G6/A6 (upper panel) and U5 (lower panel) by RbFox RRM (left: 5’-UGCAUGU; right: 5’-UGCAUAU). Side chains of key residues in the interactions (sticks) are labeled. **b**, Comparison of the two RbFox-RNA structures (blue: complex with 5’-UGCAUGU; orange: complex with 5’-UGCAUAU). **c**, Structural path for the long-range conformational rearrangement of RbFox upon binding to the non-cognate RNA (color scheme as in panel **b**; arrows: reorientation of the side chains involved in the path).

Collectively, these structural features indicate that RbFox employs structurally distinct binding modes for the cognate and the non-cognate RNA, sacrificing the binding energy of multiple hydrogen bonds and *π*-*π* stacking interactions in order to accommodate a non-cognate sequence. These observations provide structural evidence for the thermodynamic penalty model.

Equally intriguingly, the non-cognate RNA induces a distinct long-range conformational rearrangement in RbFox, which remodels the surface on the protein distant from the RNA binding site (**Fig.6b,6c**). The side chain of R118 rotates to interact with the Watson-Crick face of A6, thereby losing a hydrogen bond with the Hoogsteen face of G6, that can only be formed with the cognate RNA (**Fig.6c**). The rotation of R118 leads to a reorientation of the side chain of K117. These rearrangements, together with the reoriented side chain of N190 and flipping out of the side chain of E164, are necessary to relieve steric clash (**Fig.6c**). This domino-like effect causes the movement of the N-terminal loop, β3-α2 loop and the α2 helix, propagating the conformational change to peripheral structural elements α1 and β2 and the connecting loop. A new surface is created by the above rearrangements, exemplified by the negative-charged residue E164 flipping out and altering the protein surface electrostatics (**Extended Data Fig.9**). Thus, a single nucleotide change in the RNA triggers not just binding mode alterations at the RNA binding site, as would be expected, but also changes at a distal surface, where other factors potentially interact with RbFox to modulate its biological activities^25^.

## DISCUSSION

Wide specificity variations are critical determinants of RBP function^1,2,5,6,8,9,12^. Defining the molecular mechanisms that determine RBP specificity is thus necessary for understanding RNA biology. Here, we have shown that a highly specific RNA binding module, the RNA recognition motif of the RbFox proteins, accomplishes extraordinary sequence specificity through a previously unknown mechanism, the use of two structurally distinct binding modes. One binding mode exclusively binds cognate and near-cognate RNAs with high affinity, while the other binding mode accommodates all other RNAs, but with markedly reduced affinity.

The notion of distinct binding modes for the RRM of RbFox is based on three converging independent lines of evidence. First, the affinity distribution of RbFox for all 16,384 7-mer RNA sequence variants is bimodal (**Fig.1**). In contrast, affinity distributions for other RBPs are unimodal^6,7,10^, reflecting the gradual incremental effects of protein sequence variations on RNA affinity^20^. A single binding mode can be modeled by either a position weight matrix or more complex models considering functional coupling between bases^22,26^ (e.g., pairwise interaction matrix). However, the bimodal affinity distribution of RbFox cannot be described by a single binding model.

Second, mutations in RbFox affect the two binding modes differently. The mutations only slightly decrease the affinity for the cognate RNA variant, but markedly increase affinity for all other variants, thereby reducing overall specificity (**Fig.3**).

Third, structural differences between complexes of RbFox with the cognate and a non-cognate sequence are observed (**Fig.5,6**). Beyond expected differences at the RNA-protein interface, we also identify pronounced differences in regions distant from the RNA binding site. In other RBPs for which structures with different RNA variants have been reported, structural variations are generally concentrated at the RNA binding sites^27–29^, even where relatively large changes are introduced in the protein^28,30^. Our data with RbFox reveal instead that structural rearrangements in the RRM are communicated to a distant surface. While this is not unprecedented, we observe that the conformational change depends on whether a cognate or non-cognate RNA is bound to the protein. The structural analysis suggests that the different binding modes are accomplished in a switch-like manner: the cognate variant induces one structure while non-cognate variants another. It is possible that additional, but presumably smaller structural differences occur within the non-cognate binding mode, similar to the differences that have been observed for other RBPs bound to multiple different RNAs^27–29,31^. Structural and functionally, the propagation of the conformational rearrangements to sites distant from the RNA interface reveals that RNA can act as an allosteric effector, potentially regulating the binding of other proteins to RbFox, and altering the composition of multi-component RNP complexes, including the Large Assembly of Splicing Regulators (LASR)^32^.

The use of distinct binding modes by a single RRM to achieve high sequence specificity is, to our knowledge, a novel concept for single protein domains. Distinct binding modes have been reported for proteins with multiple domains, reflecting the differential contributions of each RRM for binding to complex multimodal RNA sites; an important and well understood advantage of their modular structure ^29,33–35^. However, these scenarios differ fundamentally from the multiple binding modes of a single RNA binding module shown here. The two binding modes of the RbFox RRM enable exceptionally high sequence selectivity by enabling the imposition of a high thermodynamic penalty on the non-cognate sequences. Mutations in the RRM diminish the ability to maintain the high thermodynamic penalty, the affinity for non-cognate variants therefore increases and specificity decreases (**Fig.3**).

We speculate that the two binding modes evolved to allow sharp discrimination between small sequence variations within a single RRM, which might be impossible to accomplish within a single binding mode, as reflected by the generally relaxed specificity of many RRMs^36^. To accomplish an RbFox-like difference between cognate and near cognate sequences within a single binding mode would require a range in the affinity distribution that might not be physically possible, biologically desirable, or both. Since RNA-protein association is universally limited by diffusion, affinities in the sub-nanomolar range or higher are necessarily associated with long lifetimes of the RNA-protein complex^37^, which might be incompatible with RBP functions in pre-mRNA splicing regulation, where remodeling of RNA-protein interactions occurs on the scale of minutes or faster^1,2,37^. Low affinities for RNA-protein interactions are limited by the electrostatic properties of RNA, which promote “non-specific” interactions with proteins that are usually in the low to mid micromolar range^1,2,37^. We therefore propose that the dual binding modes for RbFox evolved to overcome the physical limits of sequence discrimination imposed by a single binding mode.

It remains an open question why RbFox requires higher specificity than other RBPs. Recent data show that RbFox binds and promotes biological effects also at near-cognate RNA sites in cells, but only at increased cellular RbFox concentrations^20^. The extraordinary specificity of RbFox might thus be necessary for high fidelity binding to the cognate sequence at low RbFox concentrations, which is likely critical for accurate control of splicing networks in brain, heart, and muscle, and during embryonic development^14^. In addition, the different conformations of RbFox on cognate and non-cognate sequences described herein, might promote the assembly of distinct regulatory complexes, which allow RbFox to transmit even minute changes in RNA sequence to potential binding partners. The structural differences between the two binding modes thus add a previously unappreciated layer of regulation to RBP biology.

## Acknowledgements

This work was supported by the NIH (GM118088 to E.J., GM126942 to G.V.) and National Natural Science Foundation of China (31900863 to F.Y.). We thank Michael E Harris, Hsuan-Chun Lin (Univ. Florida, Gainsville, FL) and Ulf-Peter Guenther (DKMS, Dresden, Germany) for advice and technical assistance.

## Conflict of interest statement

GV is a co-founder of Ithax Pharmaceuticals and Ranar Therapeutics. EJ is a co-founder of Bainom Inc.

## Data Accessibility

Atomic coordinates and NMR restraints of the structures have been deposited in the Protein Data Bank under 7VRL.

## MATERIALS AND METHODS

### Protein expression and purification

Expression and purification of RbFox RRM and RbFox^mut^ were performed as previously described^17^. Briefly, the recombinant plasmid was transformed into BL21 (DE3) competent cells. Transformants were grown in LB media at 37°C to OD_600nm_ = 0.6. The recombinant proteins were overexpressed in E. coli at 18°C overnight upon induction by addition of 0.5 mM IPTG. The cells were harvested by centrifugation and resuspended in lysis buffer (20 mM Tris at pH8.0, 150 mM NaCl and 1 mM DTT) and lysed by sonication at 4°C. The crude extracts were centrifuged at 27,000g for 60 min to remove the cell debris. To purify the recombinant proteins, the supernatant was applied onto a HisTrap column (GE Healthcare) and a Heparin column (GE Healthcare) in succession. Protein concentrations were measured by UV absorbance at 280 nm and confirmed by Bradford assay.

Protein preparation for NMR experiments required complete removal of traces of RNase contaminants, due to the long incubation and weak binding of wt RbFox to the non-cognate RNA. To accomplish this, RbFox was double tagged with both His^6^ and GST at its N-terminus and purified with HisTrap and GSTrap columns in succession, followed by a Heparin column to remove residual non-specifically bound RNAs. Both affinity tags were then cleaved with TEV protease and removed by Nickel affinity chromatography. The purified protein eluent was concentrated and loaded onto a Superdex 75 10/300 GL column equilibrated with NMR buffer (10 mM Sodium Phosphate at pH6.0 with 30 mM NaCl). Aliquoted protein samples were flash frozen for storage at −80°C.

### RNA substrates for equilibrium binding and HiTS-Eq measurements

RNA substrates were purchased from Dharmacon (Lafayette, CO) and Sigma-Aldrich (St. Louis, MO). RNA substrates were 5’ end labeled using γ^32^P-ATP and T4 Polynucleotide Kinase (NEB, MA)^38^. Uniformly radiolabeled RNA substrates were purified with 20% denaturing PAGE (acrylamide/bis 19:1, 7M urea, 0.5 x TBE) and concentrations were quantified by scintillation counter.

For the HiTS-Eq randomized RNA pool, 5’ end radiolabeled randomized RNA substrates containing a central segment of 7 randomized (*NNNNNNN*) nucleotides flanked by illumina sequencing primers (underlined sequences) were annealed with two complementary DNA oligonucleotides (underlined sequences) (**Table 1**). The RNA-DNA complexes were purified by 15% non-denaturing PAGE (acrylamide/bis 19:1, 0.5 x TBE) and concentrations were quantified by scintillation counting^38^.

**Table 1.**
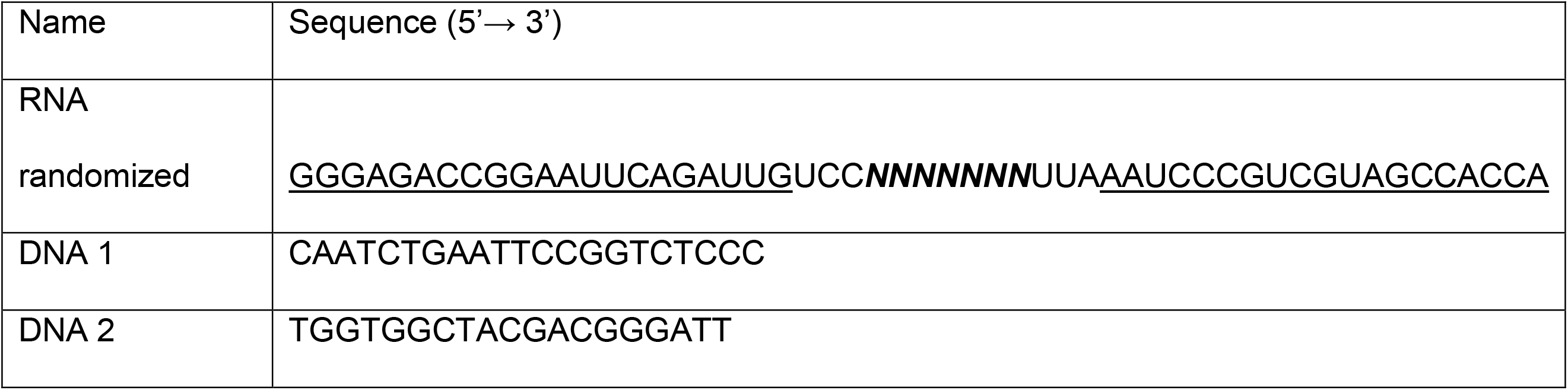
Substrates for HiTS-Eq measurements.

### Equilibrium binding with individual RNAs

RNA-protein binding reactions for wt RbFox with consensus RNA and for RbFox^mut^ with all RNAs were performed at 30 °C (10 mM Tris 7.5, 150 mM NaCl, 3.4mM EDTA, 0.001% NP-40, pH 7.5, 5’ end radiolabeled RNAs: 1 nM) and protein concentrations as indicated in the Figures (for sequences see **Table 2**). RNA and protein were incubated for 10 min. Longer incubation times did not change the observed fractions of bound RNA. Samples were then loaded on 8% non-denaturing PAGE. Gels were dried, and radioactivity in bound and free RNA was quantified using Phosphorimager (GE) and ImageQuant 5.2 software (GE Healthcare, IL). Plots of the fraction bound RNA vs. protein concentrations were fitted against the quadratic binding equation using KaleidaGraph (v3.52).

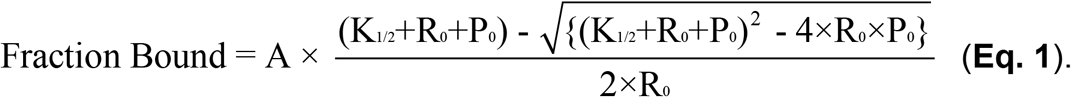

(A: reaction amplitude, K1/2: apparent dissociation constant, R0: RNA concentration, P0: protein concentration).

**Table 2.**
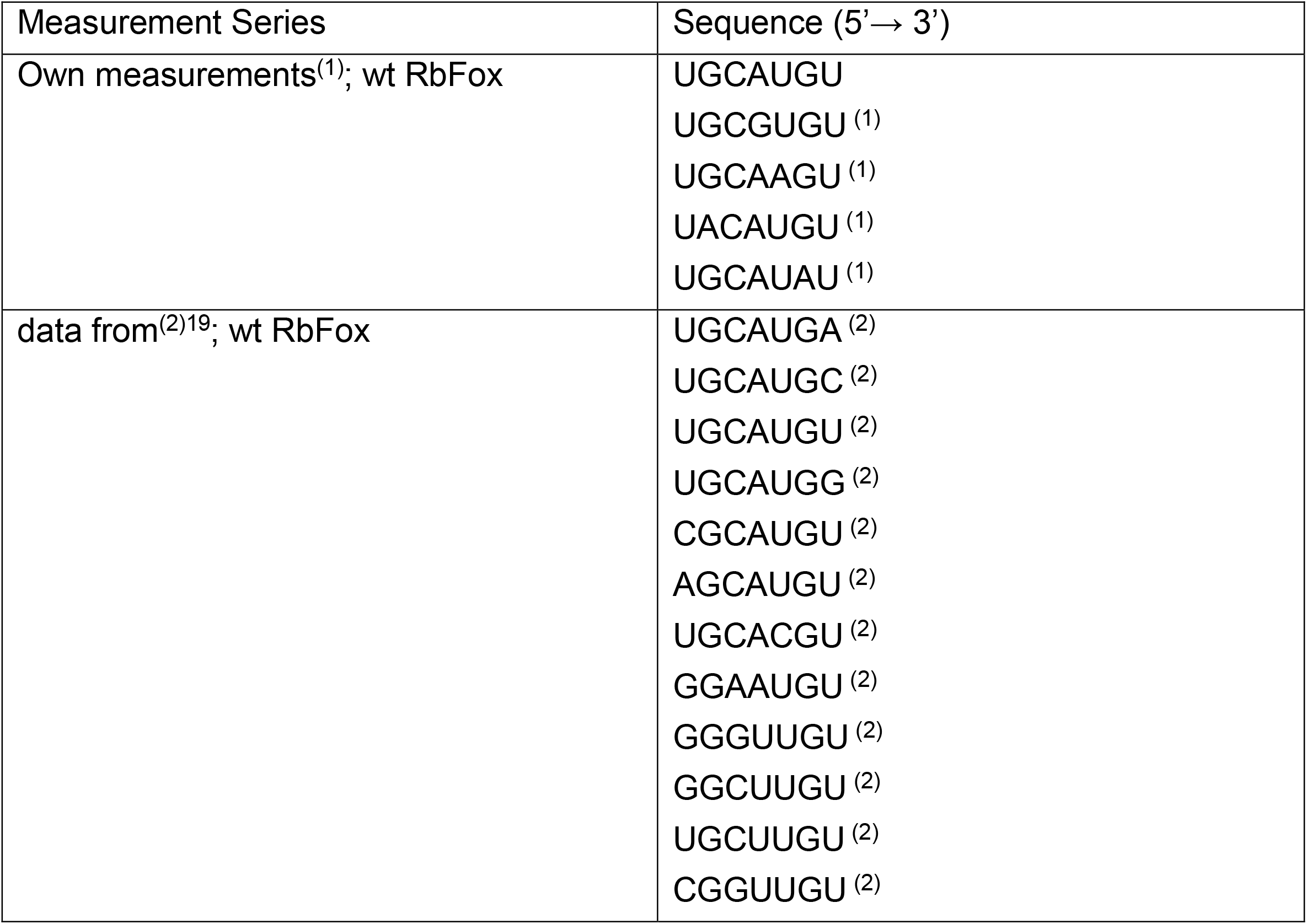
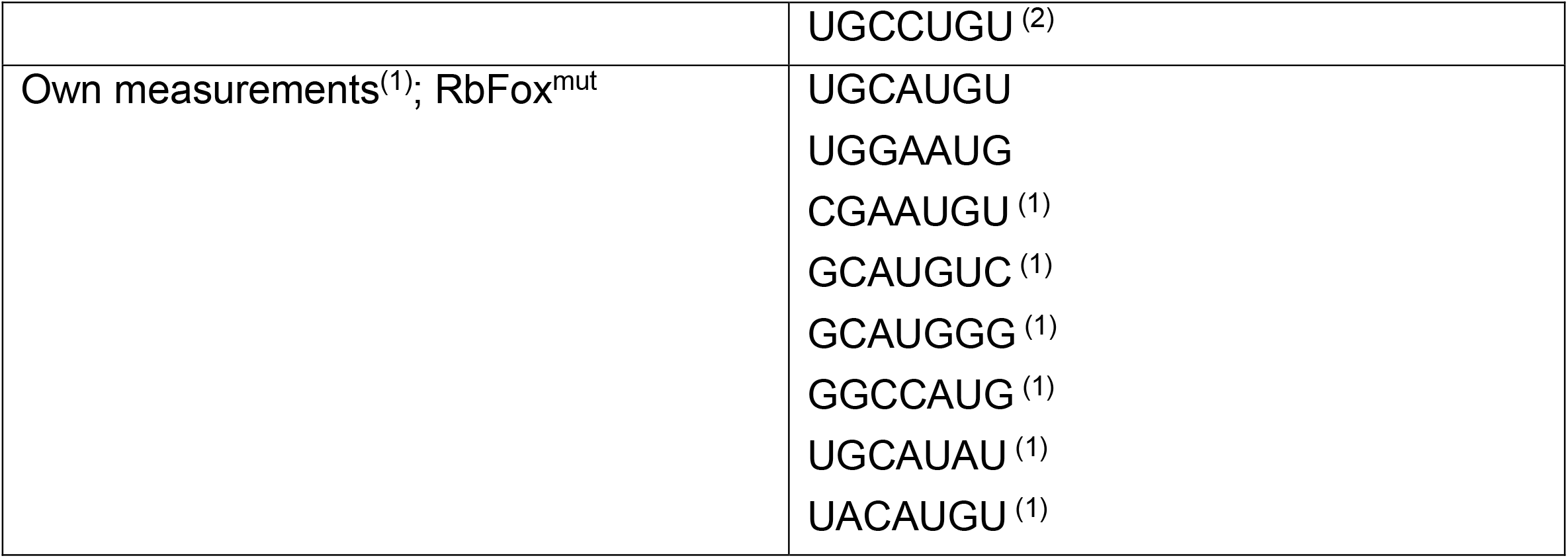
Substrates for affinity measurements with individual RNAs. ^(1)^ RNAs used in competition experiments. ^(2)^ Measurements were conducted by surface plasmon resonance (BIACORE)^19^.

Affinities for wt RbFox for non-consensus RNAs were measured by competition with consensus RNA, due to low affinities of wt RbFox for these RNA variants. Consensus RNA (final concentration 1 nM) was incubated with RbFox1 for 10 minutes. Competitor RNAs at increasing concentrations were added and incubated for 10 minutes. Longer incubation times did not change observed fractions of bound and free RNA. Samples were loaded on 8% non-denaturing PAGE, as above. Gels were dried and radioactivity in bound and free bands were quantified using a Phosphorimager (GE) and ImageQuant 5.2 software (GE Healthcare, IL). Plots of the fraction bound RNA vs. chase RNA concentrations were fitted against the binding equation for competitive inhibition using KaleidaGraph^39^.

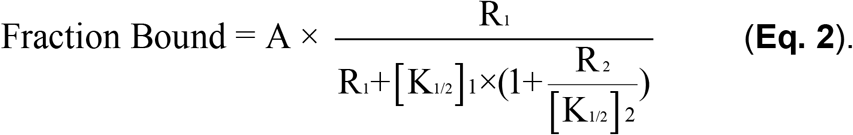

(A: reaction amplitude, R_1_: Substrate concentration, R_2_: Competitor concentration, [K_1/2_]_1_: apparent dissociation constant for substrate, [K_1/2_]_2_: apparent dissociation constant for competitor)

### HiTS-Eq measurements and sequencing library preparation

HiTS-Eq reactions (50 μL) were performed with 1 nM of 5’ end radiolabeled RNA pool with increasing protein concentrations, as indicated in the figures. RNA and protein were equilibrated for 10 min. Longer incubation times did not change the fraction of bound and free RNA. Bound and free RNA were separated by 8% non-denaturing PAGE. The gel was exposed for 30 min to autoradiography film, free RNA species were located, cut out, eluted from the gel and RNA was extracted and recovered as previously described^38^.

The HiTS-Eq libraries for Illumina next generation sequencing (NGS) were generated from the eluted RNA as previously described^7,10,11,21^. The RNA was reverse transcribed into cDNA with Superscript III Reverse Transcriptase (Invitrogen, CA). The cDNA was PCR amplified into HiTS-Eq libraries using index barcode primers (**Table 3**). The HiTS-Eq DNA libraries were purified with 8% non-denaturing PAGE. DNAs with 137 bp were cut out and extracted as previously described^7,10,21^. DNA concentration and quality were analyzed using a DNA Bioanalyzer chip (Agilent, CA). Subsequently, the HiTS-Eq libraries for different protein concentrations were pooled (equimolar) and sequenced using 50 bp single-end reads on Illumina HiSeq 2500.

**Table 3.**
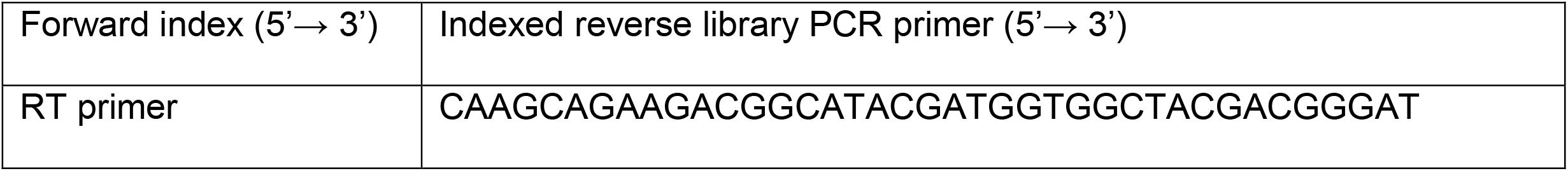
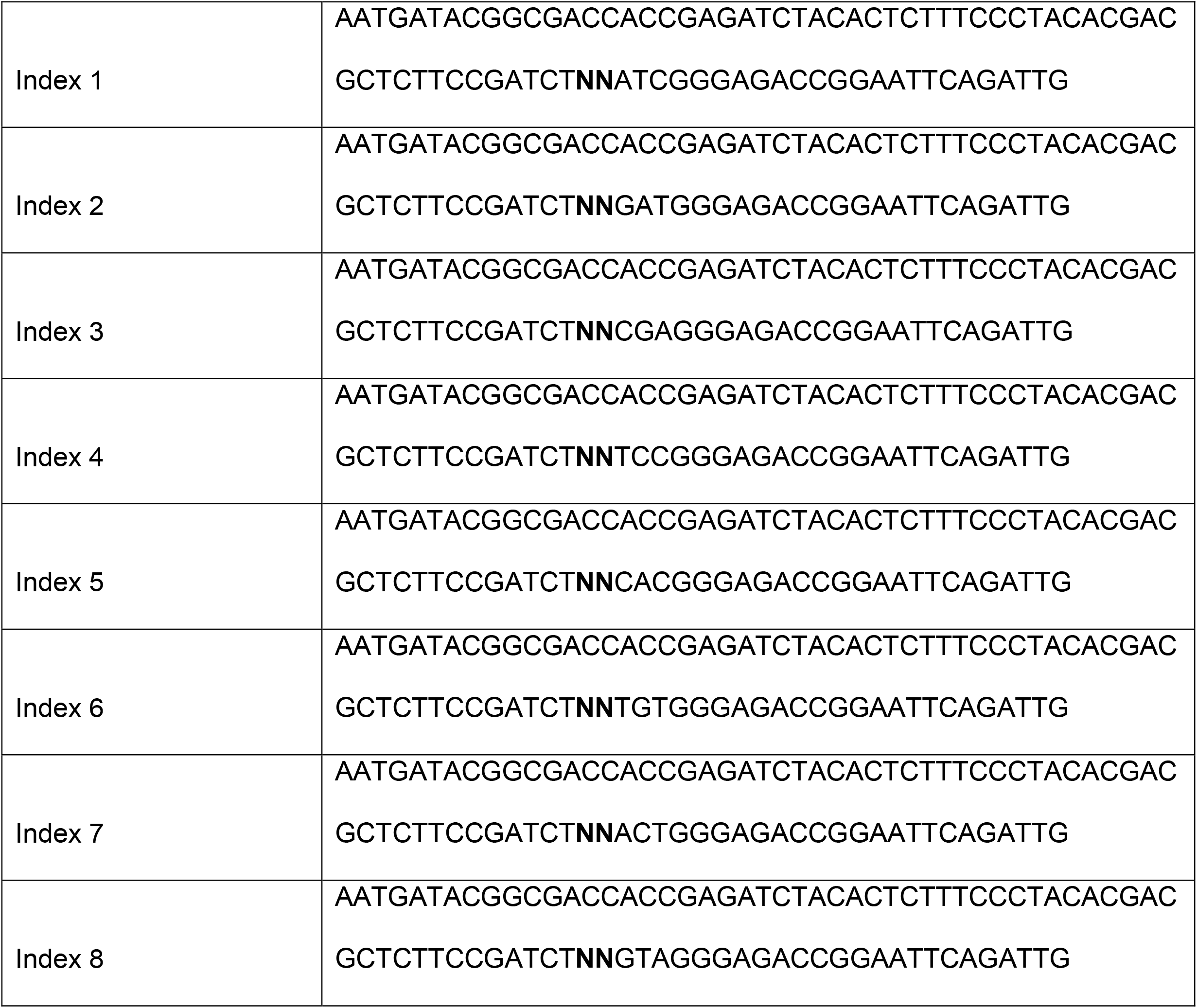
Primers for HiTS-Eq libraries. Index barcode forward primer (***NN***, degenerate nucleotides)

### Processing of Illumina sequencing data

Raw sequencing reads were quality checked with FastQC v0.11.5 and de-multiplexed based on their corresponding index barcode primers using Novocraft v3.0.8 (http://novocraft.com) or the Fastx-Toolkit v0.0.13 (http://hannonlab.cshl.edu/fastx_toolkit/). The sequencing reads were then aligned to the sequence of nucleotides 6-29 of the single strand randomized substrate, allowing one mismatch but no gaps using Perl scripts. The counts of the individual 7-mer sequences and two degenerative random nucleotides in the index barcode primers were added and exported into excel tables using Perl scripts (https://github.com/hsuanchunlin/HiTS-EQ).

### Calculation of binding constants from HiTS-Eq data

Relative equilibrium constants (*K*_A,rel_) were calculated from concentration dependent changes of the 7-mer sequence variants analogous to the procedure described previously^7,10,21^. The ratios of the two RNA substrates S_1_ and S_2_ at a given protein concentration (E) was described using a competitive binding scheme according to: S_1,0_ and S_2,0_ are initial concentrations of the RNA substrates S_1_ and S_2_ at HiTS-Eq control reaction, and equilibrium constants are described as *K*_1_ and *K*_2_.

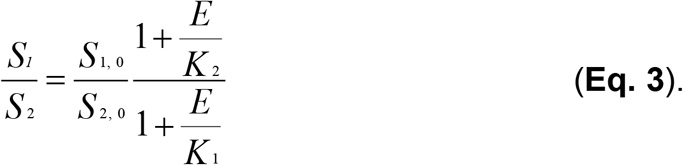

(S_1,0_ and S_2,0_: initial concentrations of the RNA substrates S_1_ and S_2_ at the HiTS-Eq control reaction, *K*_1_ and *K*_2_: equilibrium constants)

The association constant (*K*_2_) for S_2_ is:

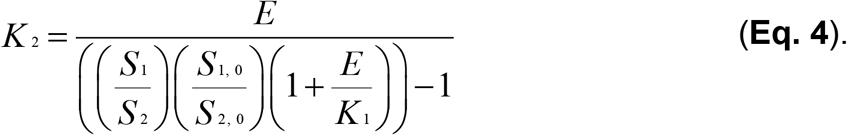

The relative association constant for a given RNA variant (*K*_2,rel_) is the value for the given RNA variant (S2), normalized by the value for substrate S1 [UGCAUGU], according to:

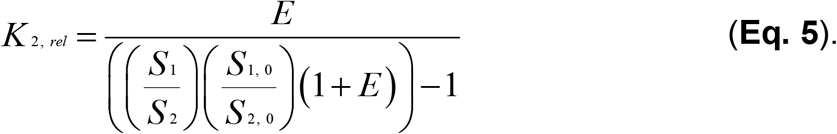

*K*_A,rel_ values for each 7-mer RNA variants were calculated using the sequencing read counts over protein concentrations. The *K*_A,rel_ values for 5-mer RNA variants were determined by splitting each 7-mer substrates into 3 non-overlapping 5-mer substrates and averaging their *K*_A,rel_ values.

### Quantitative binding models

The relative association constants were fitted with the Position Weight Matrix (PWM) model as previously described^7,10,21^. For each sequence variant, the predicted association constant (*K*_A_) value is determined by a set of linear coefficients at individual nucleotide positions, according to:

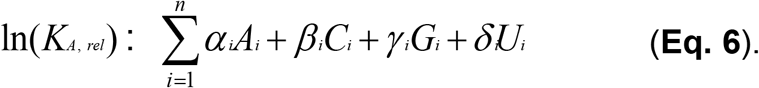

The numbers for N_i_ (N = A, C, G, U) are based on the nucleotide identity at position i (i=1 – n, n refers to the length of the sequence motif). Ni was assigned to value 1 for matched nucleotide at position i, and 0 otherwise. The cognate sequence motif 5’-UGCAUGU or 5’-GCAUG for 5-mers were used as baseline and excluded from the linear regression for the PWM models. Linear coefficients for the PWM model correspond to the parameters α_i_, β_i_, γ_i_ and δ_i_, i=1 - n.

The Pairwise Coupling (PWC) model considers all pairwise interactions between two nucleotides^7,10^. Interaction coefficients between individual nucleotide positions were added to the PWM model to fit the *K*_A,rel_ data from HiTS-Eq in equation, as previously described^7,10,21^. Interaction coefficients values (I_n_) with T value larger than 3.5 were considered statistically significant. Obtained interaction coefficients values were plotted as heatmaps.

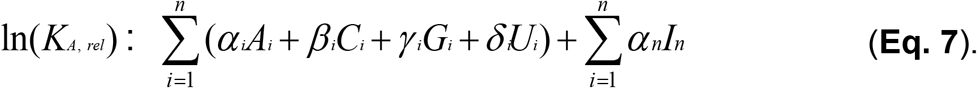

### NMR sample preparation and data collection

RNAs for NMR measurements were purchased from Integrated DNA Technologies. Oligonucleotides were dissolved in RNase-free water, desalted using a PD MiniTrap G-10 and re-suspended in NMR buffer after overnight lyophilization. ^13^C, ^15^N labeled RRM of RbFox (residues 109-208) was prepared in minimal M9 medium supplemented with 1 g/l ^15^NH_4_Cl and 2 g/l ^13^C-glucose. The complex was formed by titrating unlabeled RNA into ^13^C, ^15^N labeled protein, monitored by recording the ^15^N HSQC’s of the protein. NMR experiments were performed on Bruker Avance 600 and Avance 800 spectrometers equipped with HCN cryo-probes and pulse field gradients. NMR data were processed with NMRPipe^40^ and analyzed with CCPNMR^41^. 2D F1-, F2-filtered NOESYs^42^ (mixing times 100 ms and 300 ms) and 2D TOCSY (mixing time 80 ms) were collected on unlabeled RNA in complex with ^13^C, ^15^N labeled protein in D2O at 298 K to assign the aromatic and sugar protons of the RNA. 2D ^15^N/^13^C HSQCs, 3D HNCACB, CBCA(CO)NH, HNCO, HN(CA)CO, HBHA(CO)NH and HCCH-TOCSY spectra were collected on ^15^N, ^13^C labeled protein in complex with unlabeled RNA to assign the backbone and the non-aromatic side-chains of the protein. 2D ^13^C HSQC with carbon centered at ~ 125 ppm and 3D ^13^C NOESY-HSQC (mixing time 150 ms) were also recorded to assign aromatic side chains of the protein.

### NMR resonance assignments and structural calculation

Protein assignments followed the regular protocol using triple resonance spectra listed above. RNA resonance assignments were obtained from 2D NOESY and TOCSY by comparison with chemical shifts of 5’-GCAUG motif of pre-miR20b stem loop complexed with RbFox and those reported for 5’-UGCAUGU-RbFox complex^17^. Manually assigned intermolecular NOE distance restraints derived from 3D NOESYs at 100 ms mixing time were separated into three ranges based on the cross-peak intensities, strong (1.8 Å −3.5 Å), medium (1.8 Å-4.5 Å) and weak (1.8 Å-5.5 Å). Additional NOEs observed only in 2D NOESY with 300 ms of mixing time were assigned as very weak (1.8 Å-6.5 Å).

The NOE distance restraints for the complex could be divided into two parts, ‘experimental’ NOE derived experimentally, as described below and ‘virtual’ NOE, predicted from the miR20b-RbFox complex based on chemical shift similarities. The ‘experimental’ NOE list was obtained from the combination of: intra-protein NOEs automatically assigned from ^15^N and ^13^C NOESY-HSQCs obtained by CYANA^43^; intra-RNA NOEs manually assigned from 2D F1-, F2-filtered NOESYs spectra; and intermolecular NOEs manually assigned from 2D F1-filtered, F2-edited NOESY and 3D ^13^C F1-filtered, F3-edited NOESY-HSQC.

Dihedral angle restraints for the conformation of sugar rings (C2’-endo or C3’-endo) were added, based on H1’-H2’ cross-peak intensities in 2D TOCSY spectrum. With all of the restraints above, 100 initial structures were generated in CYANA^43^, and the 20 structures with the lowest target function were further regularized in implicit solvent model using the SANDER module of Amber 14.0^44^. We have used ff99bsc0xOL3 force field for RNA and ff14SB force field for protein. The script for the restrained simulated annealing protocol was modified from Tolbert *et al*^45^. Protein torsion angles were obtained by TALOS+^46^. For the complex, we heated the system to 1500 K during Amber simulated annealing refinement. The 20 lowest-energy structures were analyzed with PROCHECK NMR^47^.

## FIGURE CAPTIONS, EXTENDED DATA

**Extended Data Figure 1.**
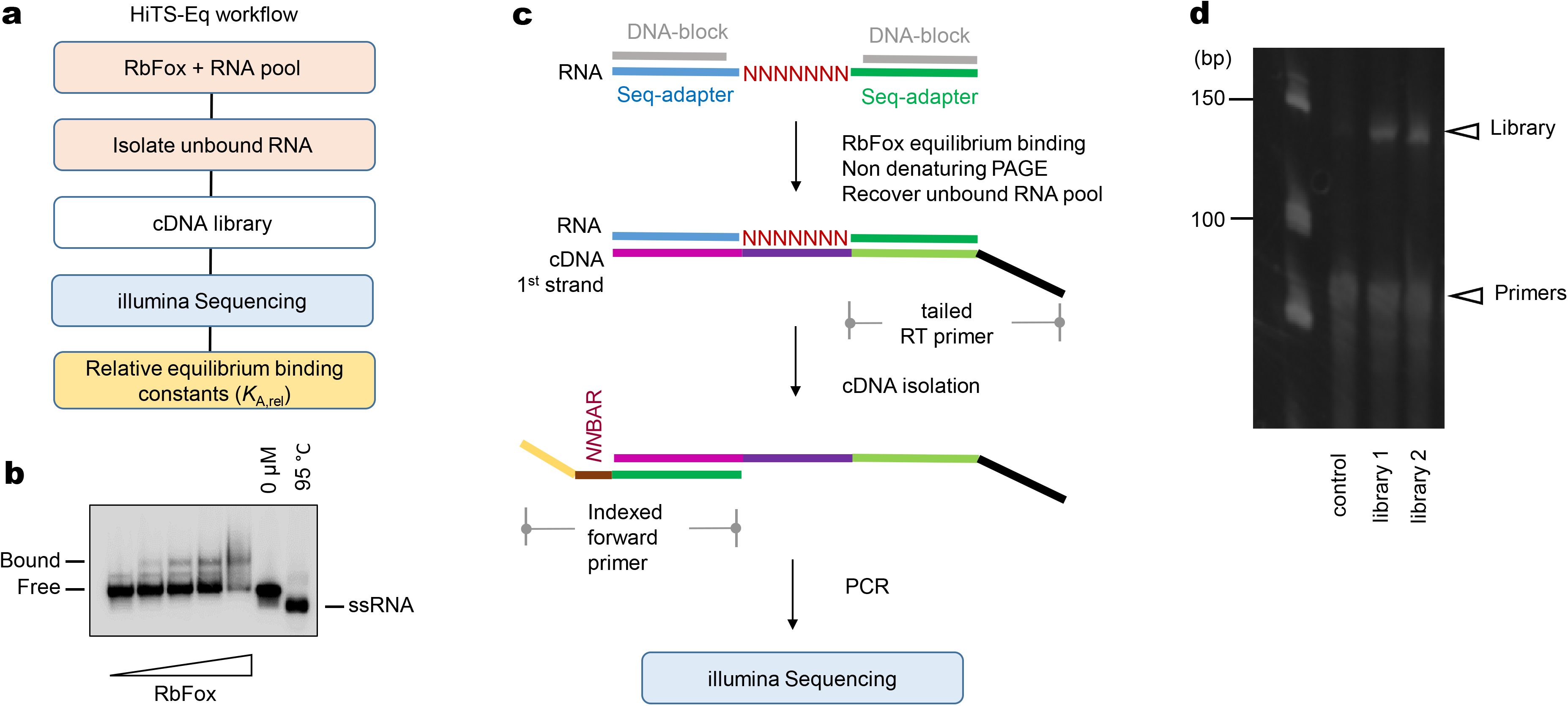
HiTS-Eq approach. **a**, HiTS-Eq workflow. **b**, Representative PAGEs for RbFox binding to the randomized RNA pool. The unbound RNA species were recovered and converted into NGS library according to the HiTS-Eq protocol. **c**, Preparation of HiTS-Eq libraries for illumina sequencing (detailed information: Materials and Methods). **d**, Representative PAGE of the HiTS-Eq libraries (expected length: 137 bp).

**Extended Data Figure 2.**
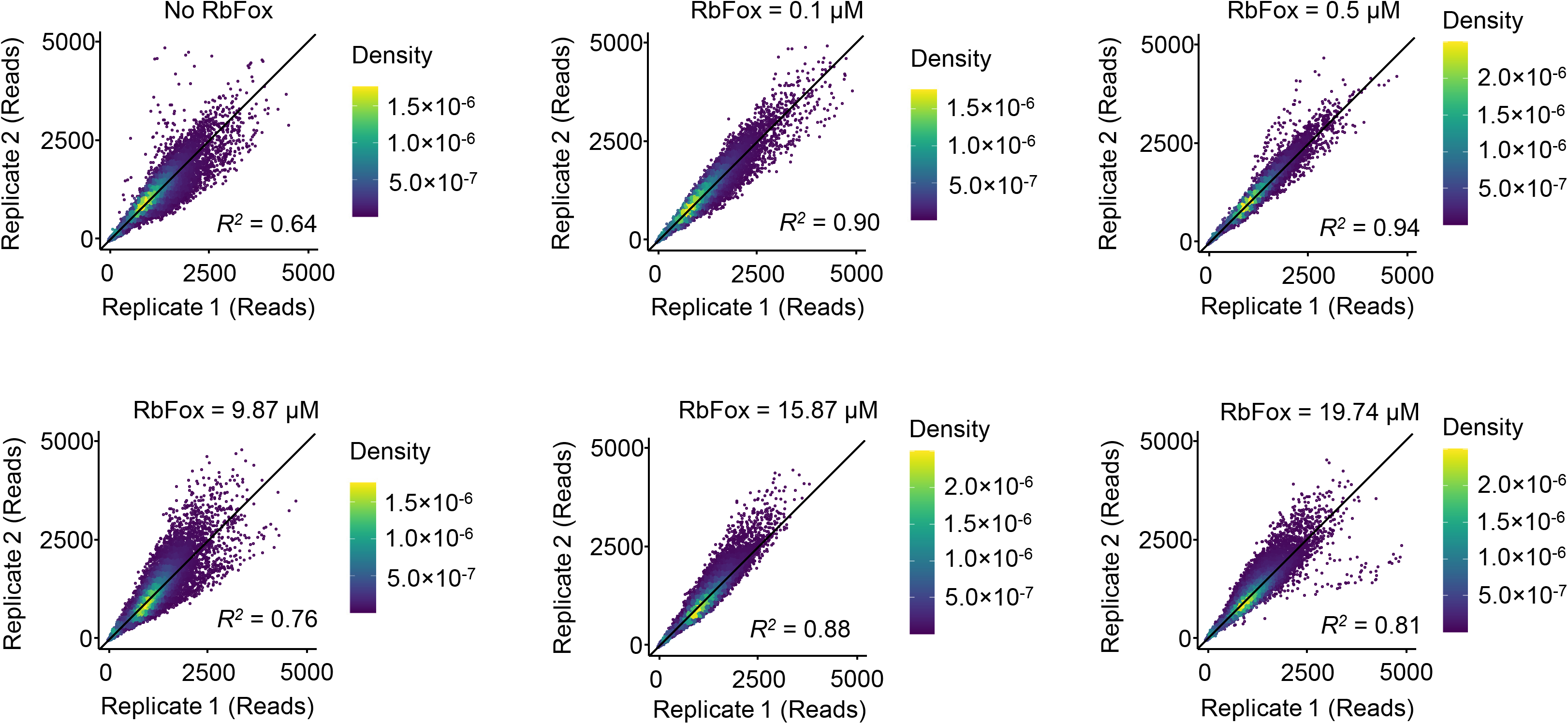
Correlation between HiTS-Eq replicates. Correlation between the two matched replicates for all 16,384 7-mer RNA sequence variants at different RbFox concentrations (line: diagonal, y = x; *R^2^*, correlation coefficient).

**Extended Data Figure 3.**
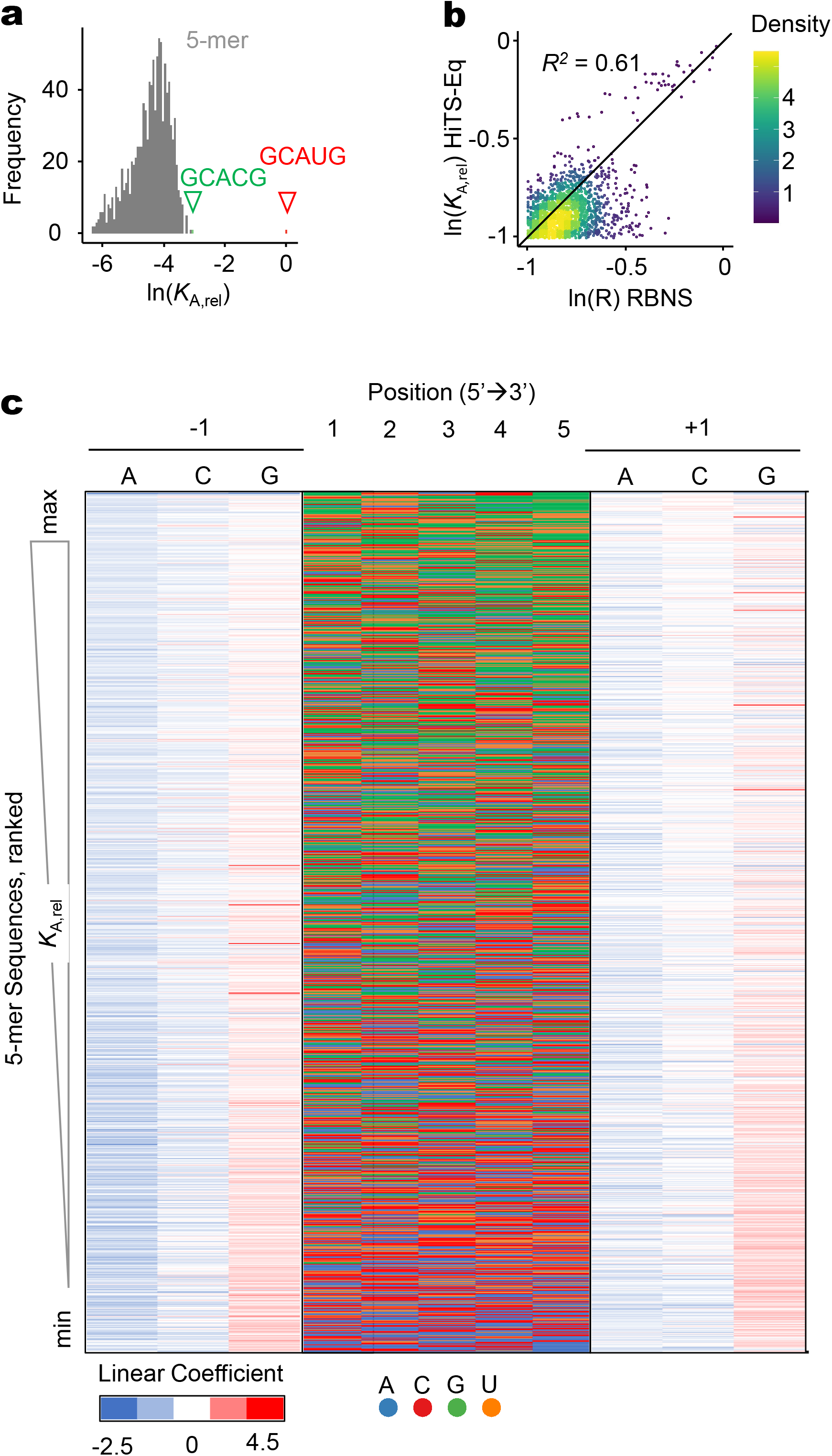
RbFox binding to 5-mer RNA variants. **a**, Distribution of relative association constants (*K*_A,rel_) of RbFox for all 1,024 5-mer RNA sequence variants (red triangle: reference 5’-GCAUG; green triangle: 5’-GCACG). **b**, Correlation between relative association constants (*K*_A,rel_) from the HiTS-Eq measurements and R value from RNA Bind-n-Seq (RBNS) measurements^16^ for all 1,024 5-mer RNA sequence variants. (line: diagonal, y = x; *R^2^*, correlation coefficient). **c**, Linear coefficient for −1 and +1 nucleotide position of all 1,024 5-mer RNA sequence variants calculated with the PWM binding model (negative values: destabilization).

**Extended Data Figure 4.**
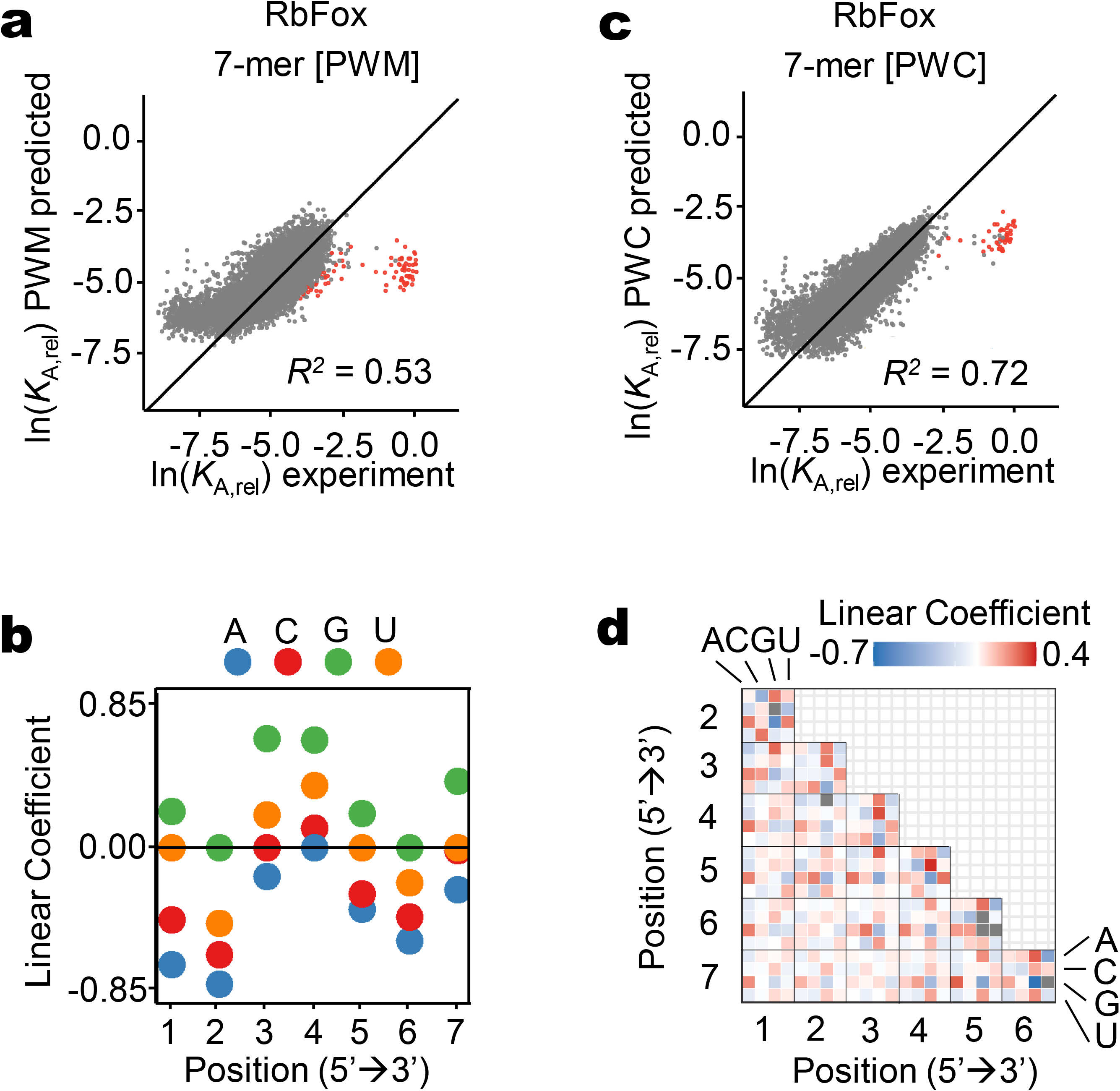
Analysis of RbFox bonding to 7-mer RNA variants with quantitative binding models. **a**, Correlation between experimental *K*_A,rel_ values for RbFox for each 7-mer with values calculated with the PWM binding model; (red dots: consensus 5-mer; line: diagonal, y = x; *R^2^*: correlation coefficient). **b**, Linear coefficient for each nucleotide position calculated with the PWM binding model (negative values: destabilization). **c**, Correlation between experimental *K*_A,rel_ values for RbFox^mut^ for each 7-mer with values calculated with the PWC binding model (red dots: consensus 5-mer; line: diagonal, y = x; *R^2^*: correlation coefficient). **d**, Linear coefficients for each pairwise coupling between all nucleotides calculated with the PWC model; (black frames: couplings in consensus 7-mer, negative values: destabilization).

**Extended Data Figure 5.**
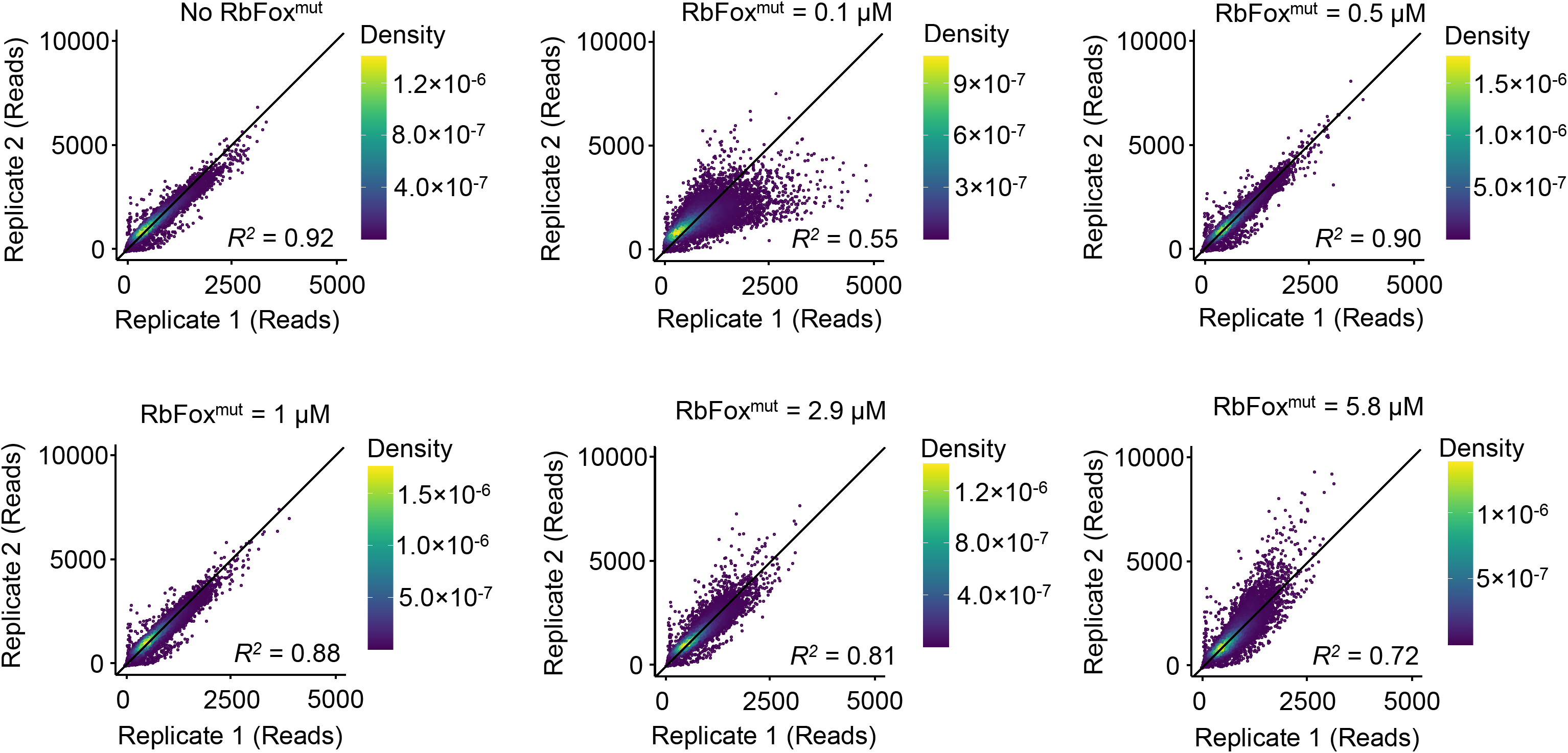
Correlation between HiTS-Eq replicates for RbFox^mut^. Correlation between the two matched replicates for all 16,384 7-mer RNA sequence variants at different RbFox^mut^ concentrations; (line: diagonal, y = x; *R^2^*, correlation coefficient).

**Extended Data Figure 6.**
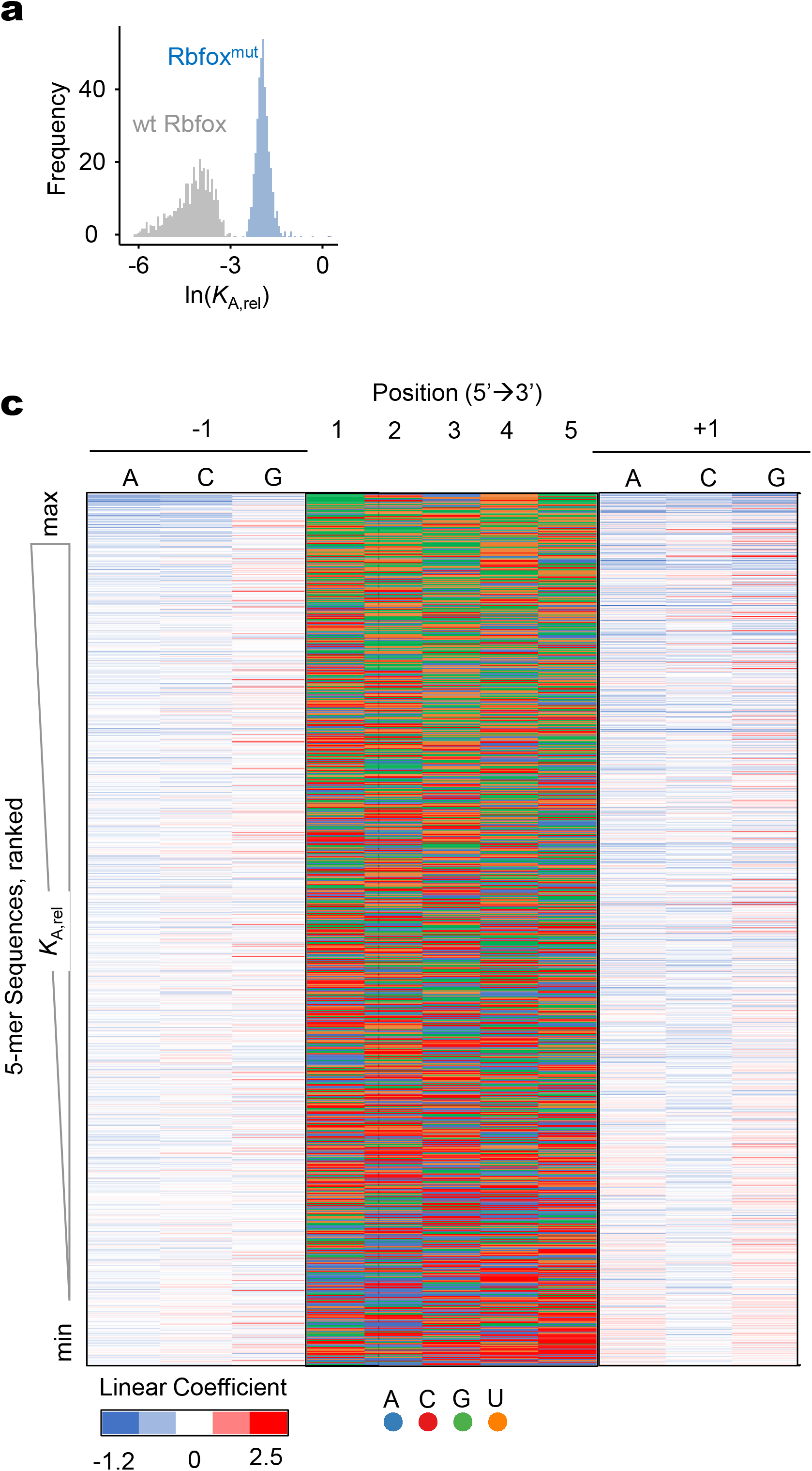
RbFox^mut^ binding to 5-mer RNA variants. **a**, Affinity distribution (*K*_A,rel_) of RbFox^mut^ for all 5-mer RNA sequence variants (blue); (bin size: 100). For reference, the affinity distribution of wild type RbFox (grey) is provided as well. **b**, Linear coefficient for −1 and +1 nucleotide position of all 1,024 5-mer RNA sequence variants calculated with the PWM binding model (negative values: destabilization).

**Extended Data Figure 7.**
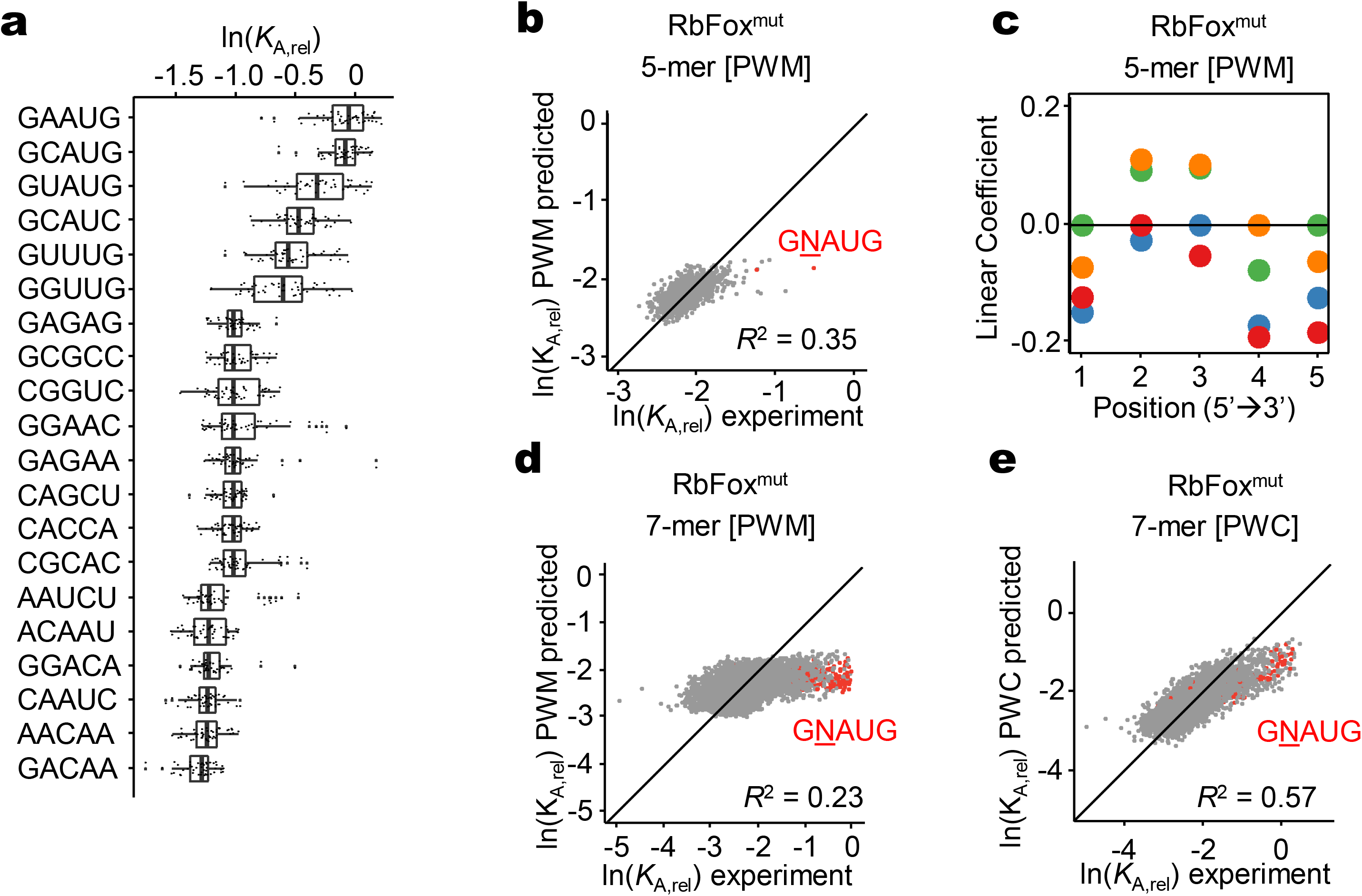
Analysis of RbFox^mut^ RNA binding with quantitative binding models. **a**, Relative affinities (*K*_A,rel_) for selected 5-mer RNA variants, as indicated on the left. 48 *K*_A,rel_ values correspond to each 5-mer; (vertical line: median; box: variability through lower quartile and upper quartile; whiskers: variability outside the lower and upper quartiles). **b**, Correlation between experimental *K*_A,rel_ values for each 5-mer (median value, panel **a**) and values calculated with the PWM binding model (red dots: consensus 5-mer; line: diagonal, y = x; *R^2^*: correlation coefficient). **c**, Linear coefficients for each nucleotide position calculated with the PWM binding model (negative values: destabilization). **d**, Correlation between experimental *K*_A,rel_ values for each 7-mer (median value, panel **a**) and values calculated with the PWM binding model (red dots: consensus 5-mer; line: diagonal, y = x; *R^2^*: correlation coefficient). **e**, Correlation between experimental *K*_A,rel_ values for each 7-mer (median value, panel **a**) and values calculated with the PWC binding model (red dots: consensus 5-mer; line: diagonal, y = x; *R^2^*: correlation coefficient).

**Extended Data Figure 8.**
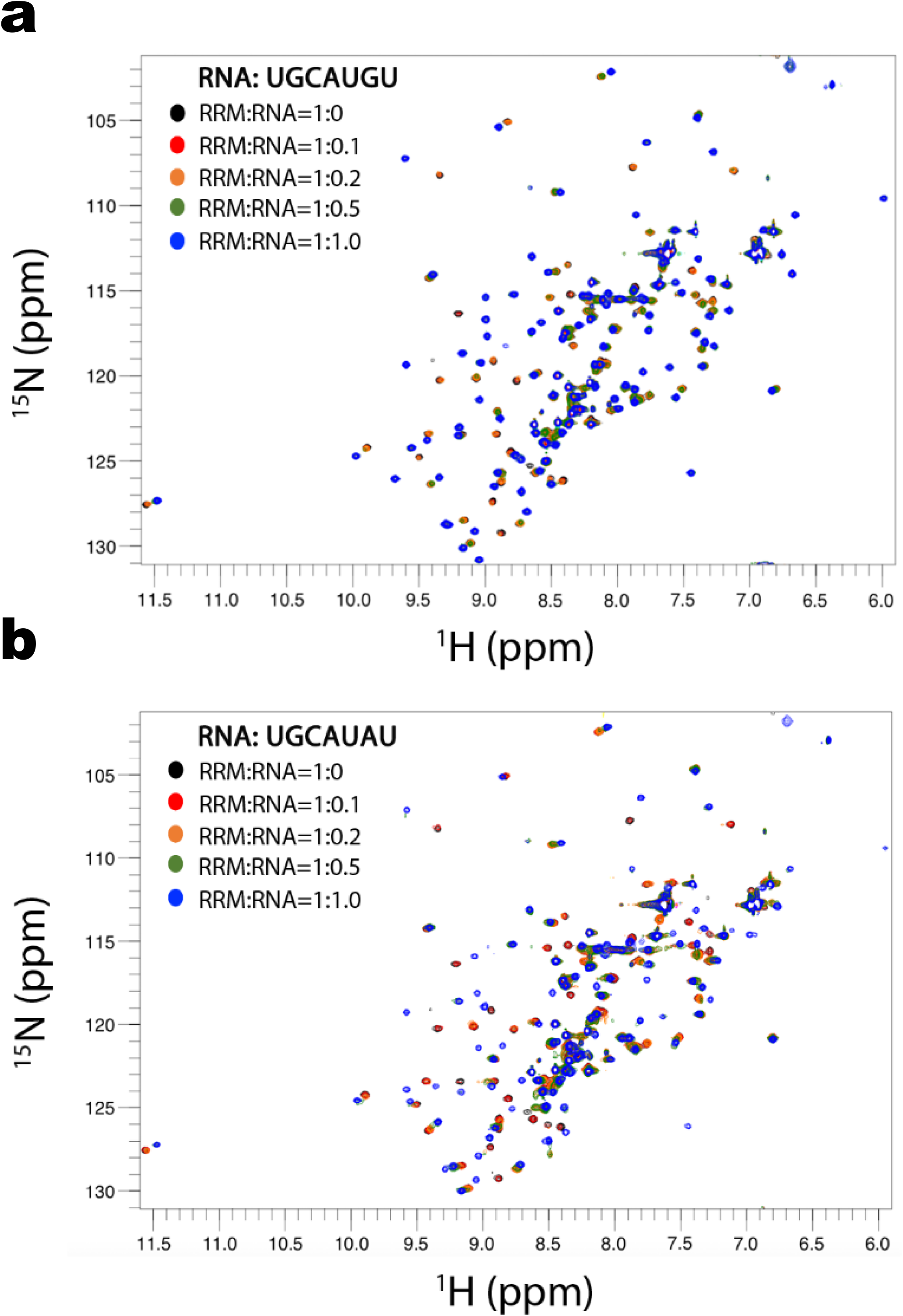
^1^H-^15^N HSQC titrations of RbFox RRM with two RNAs 5’-UGCAUGU and 5’-UGCAUAU. **a**, Superposition of ^1^H -^15^N HSQC spectra obtained with ^15^N-RbFox RRM and increasing amount of 5’-UGCAUGU RNA. The peaks corresponding to the free and RNA-bound RRMs (RRM:RNA ratios of 1:0, 1:0.1, 1:0.2, 1:0.5 and 1:1) are colored as black, red, orange, green and blue, respectively. **b**, Superposition of ^1^H -^15^N HSQC spectra obtained with ^15^N-RbFox RRM and increasing amount of 5’-UGCAUAU RNA. The color scheme is the same as in **a**.

**Extended Data Figure 9.**
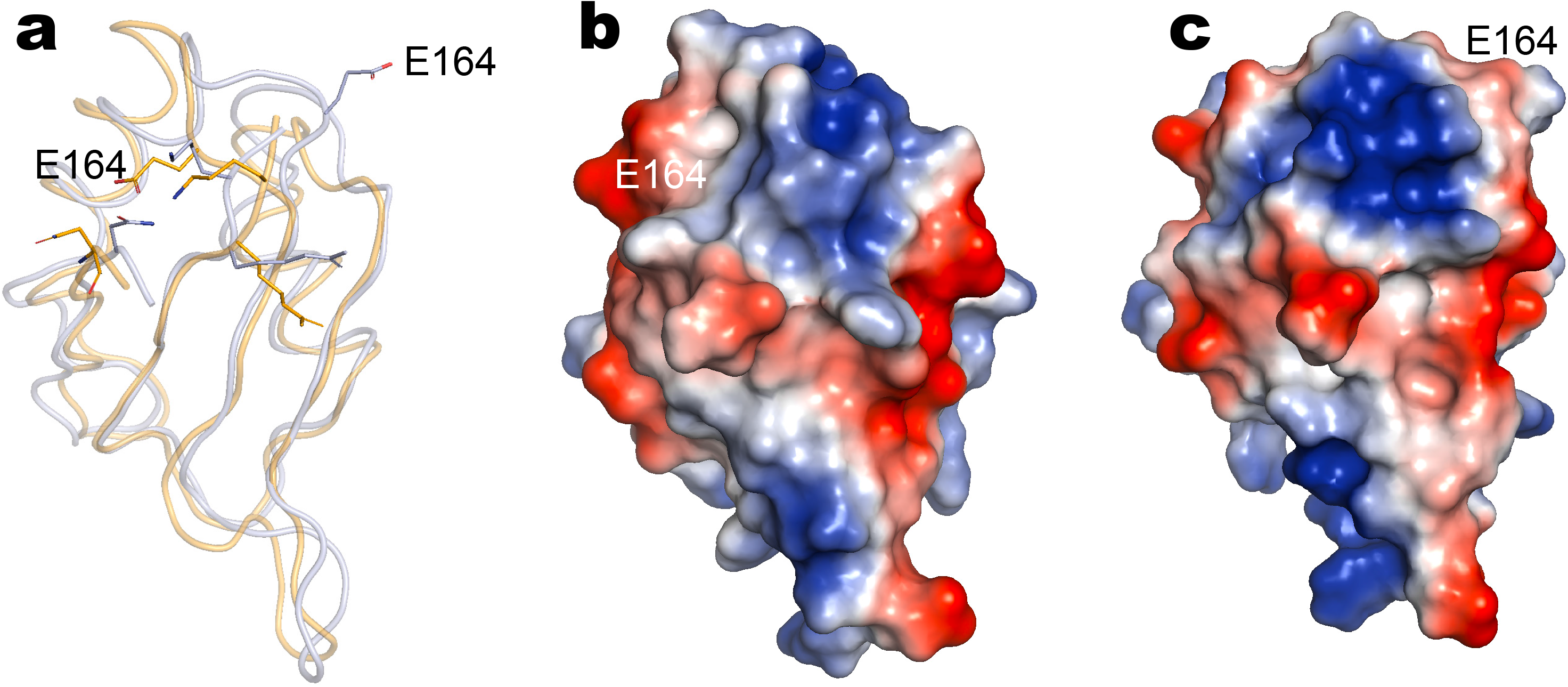
Comparison of the surface electrostatics of the two RbFox-RNA complexes. **a**, Ribbon diagrams of the superposed RbFox structures bound to the different RNA (blue: complex with 5’-UGCAUGU; orange: complex with 5’-UGCAUAU). The side chain of the flipped out E164 residue which alters surface electrostatics through its relocation is explicitly labeled. **b**, **c**, Surface electrostatics for the two RbFox-RNA complexes with 5’-UGCAUGU and 5’-UGCAUAU, respectively.

---

**Extended Data Table 1.** NMR structure statistics for Rbfox RRM bound to UGCAUAU.

| **NMR constraints** |  |
| --- | --- |
| **Distance constraints** |  |
| Total NOEs | 2020 |
| Protein-RNA intermolecular | 130 |
| RNA intramolecular | 95 |
| Protein intramolecular | 1795 |
| Protein intra-residue | 421 |
| Sequential ($\vert i-j\vert =1$) | 480 |
| Medium range ($1<\vert i-j\vert <5$) | 287 |
| Long range ($\vert i-j\vert \geq 5$) | 607 |
| Hydrogen-bond constraints | 16 |
| Torsion angle constraints | 100 |
| **Structure statistics (20 structures of lowest energy)** |  |
| Violations |  |
| NOE violations > 0.3 Å | 0 |
| Torsion angle violations > 5° | 0 |
| Ramachandran plot statistics |  |
| Residues in most favored regions | 82.0% |
| Residues in additional allowed regions | 14.6% |
| Residues in generously allowed regions | 2.2% |
| Residues in disallowed regions | 1.1% |
| RMS deviations from the mean structure |  |
| Protein backbone (Pro116-Arg194) | 0.74 |
| Protein heavy atoms (Pro116-Arg194) | 1.20 |
| RNA heavy atoms (G2-A6) | 2.70 |
| Complex heavy atoms (G2-A6 and Pro116-Arg194 ) | 2.79 |

